# Queuosine promotes *wecB*-dependent phage resistance and biofilm formation in the marine bacterium *Shewanella glacialimarina*

**DOI:** 10.64898/2026.03.05.709803

**Authors:** Pavlina Gregorova, Minna-Maria K. Heinonen, Nina Sipari, L. Peter Sarin

## Abstract

Transfer RNA (tRNA) modifications critically fine-tune translational accuracy and efficiency, influencing bacterial adaptation to environmental challenges. Among these, the queuosine (Q) modification has recently emerged as a regulator of biofilm formation, yet its role during phage infection remains unknown. Here, we investigate how Q modification links host translational control to phage infection. We show that phage infection activates the Q biosynthetic pathway, leading to elevated Q levels and enhanced translation of NAT-biased genes. This shift drives two interconnected outcomes, namely increased biofilm formation and enhanced mutagenesis mediated by translesion synthesis polymerases. We further identify conserved, slippage-prone regions within surface-related genes that act as hotspots for adaptive variation. Together, our findings uncover a novel mechanistic link between tRNA modification and phage-driven bacterial diversification.

**AUTHOR SUMMARY:** To survive changing environmental conditions, bacteria constantly adjust their metabolic processes, including how they translate their genetic information into proteins. One such adjustment mechanism involves carefully balancing the level of chemical modification on transfer RNA (tRNA), the key adapter molecule that carries amino acids but also regulates protein synthesis. In this study, we explore how a cold-active bacterium relies on queuosine, a wobble position tRNA modification, to mediate host cell responses to bacteriophage infection. We observe that as viral infection progresses, the bacterial host cells increase the level of queuosine modification present on tRNAs. This alters the efficiency by which specific proteins are produced, favoring those that are involved in biofilm synthesis and thereby form protective communities that help bacteria survive stress. At the same time, queuosine also promotes error-prone DNA replication processes that lead to an overall increase in bacterial mutation rates. Taken together, our results reveal how a small molecular change in tRNA can reshape bacterial responses to viral infection, ultimately driving genetic diversity and survival.

## INTRODUCTION

Transfer RNAs (tRNAs) are adorned with chemical modifications that play a critical role in fine-tuning translational efficiency and accuracy by modulating codon–anticodon pairing and maintaining reading-frame fidelity^1,2^. Many tRNA modifications are dynamically regulated and contribute to bacterial adaptation under environmental stresses, such as antibiotics^3–5^, oxidative stress^6–9^, temperature shifts^5,10–12^, and osmotic pressure^5,9^.

More recently, tRNA modifications have been linked to biofilm formation. Specifically, the queuosine (Q) modification has emerged as an important regulator of aggregation and biofilm development. Deletions of genes involved in Q biosynthesis have been shown to enhance bacterial aggregation and biofilm formation^13^. The Q modification is present on tRNAs with GUN codons (tRNA^Asp^_GUC_, tRNA^His^_GUG_, tRNA^Asn^_GUU_, and tRNA^Tyr^_GUA_) and the Q biosynthetic pathway appears to be at least partially conserved in most known bacteria^14–16^.

Bacterial biofilms are complex, multicellular communities formed in response to diverse environmental cues^17^. Extracellular DNA (eDNA) is one of the key structural components of biofilms and plays an important role in the initial attachment of bacteria to surfaces^18,19^. eDNA release is often mediated by quorum sensing and can occur via multiple mechanisms, including cell lysis and autolysis, active secretion via vesicles or through type IV or VI secretion systems^20,21^. Importantly, prophage activation^22,23^ or phage infection^24,25^ represent one of the predominant causes of cell lysis. While lytic phages have been mostly explored as agents to disrupt and disperse biofilms^26,27^, several studies have demonstrated that biofilm formation can also serve as a defense against phage infection^28–30^. Collectively, these observations indicate that phage effects on biofilm formation are context-dependent, shaped by the specific bacteria-phage interaction and environmental factors.

In addition to effects on biofilm formation, bacterial exposure to phage infection typically triggers the SOS response, a canonical pathway associated with DNA damage and mutagenic repair^31–33^. During the SOS response, induction of error-prone translesion synthesis (TLS) polymerases supports DNA repair yet simultaneously elevates mutation rates, driving the emergence of phage-resistant variants^34,35^. This mutagenic response may further influence the evolutionary dynamics of bacteria-phage interactions.

In this study, we elucidate the mechanisms by which Q modification regulates biofilm formation and the emergence of phage resistance during bacteriophage infection using the marine bacterium *Shewanella glacialimarina* TZS-4_T_. We demonstrate that phage infection triggers an increase in host Q levels via *queG*, leading to two interconnected outcomes driven by enhanced translation of NAT-biased genes: elevated biofilm formation and elevated mutagenesis via translesion synthesis (TLS) polymerases. Furthermore, we identify conserved, slippage mutation-prone regions within surface-related genes that serve as hotspots for adaptive variation. Notably, *S. glacialimarina* lacks NAT-decoding tRNAs and therefore depends on the high decoding efficiency of NAC tRNAs, linking its phage response and adaptive capacity to Q-mediated translational control.

## RESULTS

### Phage 1/4 and 1/40 infections induce biofilm formation of *S. glacialimarina* via NAT codon biased genes

To investigate the relationship between genes with Q codon bias and biofilm formation during phage infection, we utilized the marine bacterium *S. glacialimarina* TZS-4_T_^36^ and its two closely related phages, phage 1/4 and phage 1/40, which share genomic similarity but exhibit distinct infection characteristics^37,38^. While phage 1/4 infects *S. glacialimarina* efficiently^39^, phage 1/40 causes a milder infection (Suppl. Fig. 1a) and produces fewer nascent virions (Suppl. Fig. 1b). First, we set out to determine whether biofilm production is induced by phage 1/4 or 1/40 infection. Given the difference in infectivity, we measured biofilm production across a range of multiplicities of infection (MOI) and observed that the host forms biofilm in a phage dose-dependent manner (Fig. 1a). As expected, phage 1/4 triggers biofilm formation at a significantly lower MOI value than phage 1/40 (Fig. 1a). This is consistent with the lesser infectivity of phage 1/40 (Suppl. Fig. 1a) and confirms that partial cell lysis is required for biofilm formation.

**Figure 1.**
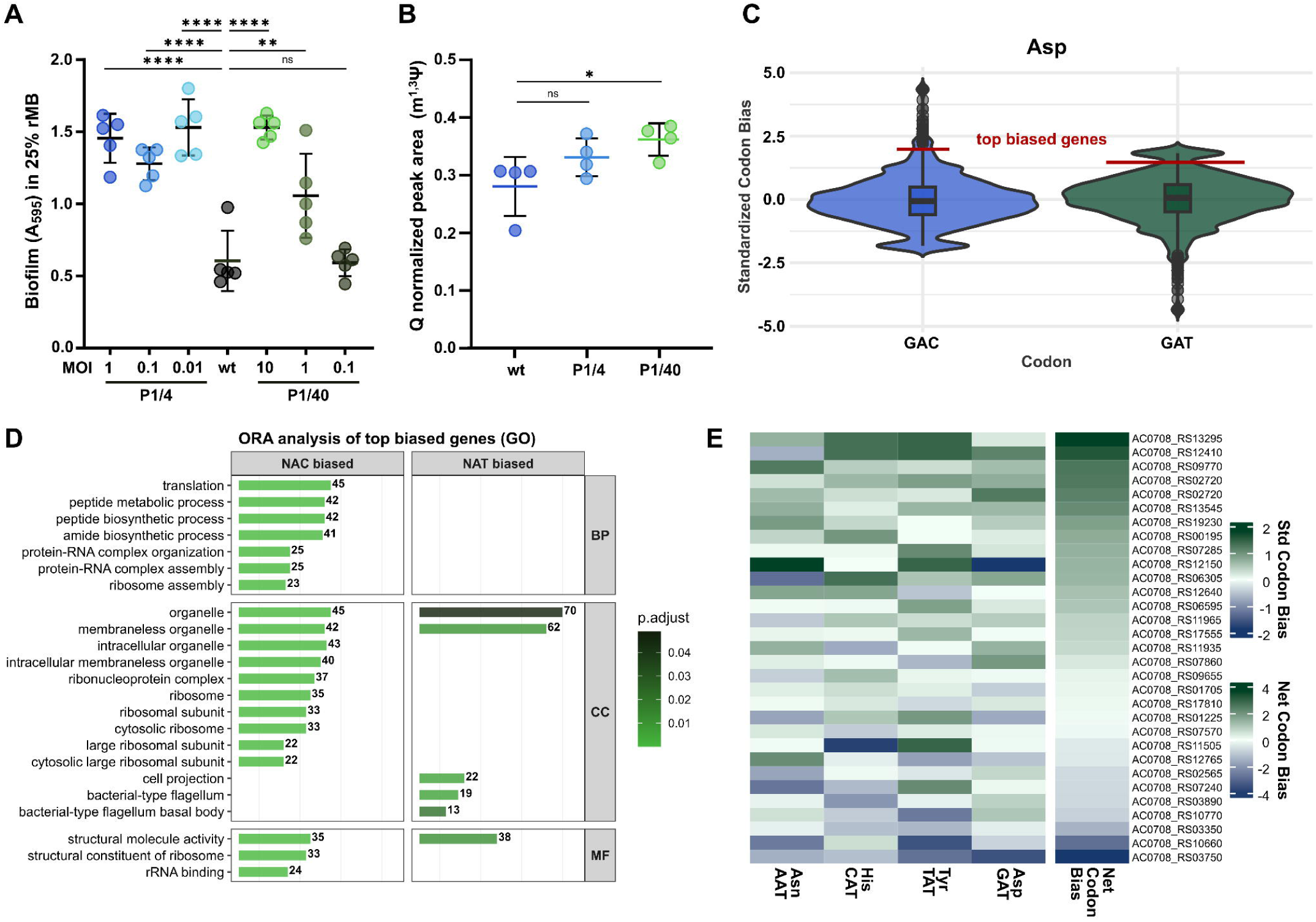
Phage induced biofilm formation is mediated by NAT codon biased genes in Shewanella glacialimarina. **a.** Phage induced biofilm measurement in 25% rMB at 24h in *S. glacialimarina* TZS-4_T_. Control without phage is labeled wt. Ordinary one-way ANOVA with Šidák’s multiple-comparisons correction (n = 5, ** p < 0.01, **** p < 0.0001, ns p > 0.05). **b.** Quantification of queuosine by LC-MS in tRNA of *S. glacialimarina* infected samples and uninfected control. Peak area is normalized to non-natural 1,3-methylpseudouridine. Kruskal-Wallis test with Dunn’s multiple-comparisons correction (n = 4, * p < 0. 05, ns p > 0.05). **c.** Example of threshold for identification of top codon biased genes for aspartic acid codon biases. Violin plots depict standardized codon bias distribution across all coding genes. Red line indicates threshold for each codon. **d.** Over-representation analysis of top biased genes for NAC (322 genes) and NAT (1302 genes) codons using GO terms. Top significant GO terms (adjusted p < 0.05, Benjamini-Hochberg correction) for Biological Process (BP), Cellular Component (CC), and Molecular Function (MF) ontologies. Terms ranked by adjusted p-value. **e.** Heatmap of standardized codon bias for each of Q decoded NAT codons in biofilm-associated genes (n = 31). Net codon bias represents the sum of individual codon biases across all codons. List of used genes is in Supplementary Table 1.

We have previously reported that host cell Q modification levels increase at the late stages of phage 1/4 infection^39^. Consequently, we (re-)examined Q modification levels for phage 1/4 and 1/40 infected cells at 70 min post-infection (p.i.), i.e., a late infection stage prior to host cell lysis (Suppl. Fig. 1b). Surprisingly, we found that phage 1/40 displays an even stronger increase in Q levels on tRNA than phage 1/4 (Fig. 1b), which suggests that upregulation of Q may constitute a general host response to phage infection rather than a measure of infection efficiency.

Q has been reported to influence the decoding of NAT/NAC codons in an organism-dependent manner^3,40,41^. To identify genes that are most affected by Q, we calculated the NAT/NAC standardized codon bias across all coding genes (Suppl. Fig. 1c), identified top codon biased genes (Fig. 1c; Suppl. Fig. 2; see Methods), and examined the functional categories of these genes (Fig. 1d). While NAC-biased genes were primarily linked to translation regulation, NAT-biased genes were enriched for bacterial flagellum and motility (Fig. 1d, Suppl. Fig. 1d).

Next, we focused on NAT/NAC codon containing genes that are annotated as biofilm forming (n = 31; Supplementary Table 1). We found that most of these genes exhibit a net codon bias towards NAT rather than NAC codons (Fig. 1e). Expanding this analysis to other genes previously associated with biofilm formation^42,43^—such as motility, secretion systems, flagella, chemotaxis, and lipoproteins (n = 353; Supplementary Table 2)—revealed a similar overall bias toward NAT codons (Suppl. Fig. 1e). These findings suggest that Q facilitates the efficient translation of biofilm-related genes and thereby contributes to biofilm induction during phage infection.

### Phage infection induces *queG* expression changes

To better understand the role of Q pathway genes in the host response to phage infection (Fig. 2a), we performed RNA-seq on infected and uninfected cultures. Among the Q pathway genes, only *queG* was upregulated (Fig. 2b; Suppl. Fig. 3a-b), suggesting that the conversion of epoxyqueuosine (oQ) to Q serves as a key regulatory step. Consistent with this, our LC-MS analysis revealed decreased oQ levels alongside increased Q levels (Fig. 2c, Fig. 1b), further supporting that *queG* regulates Q modification levels during infection.

**Figure 2.**
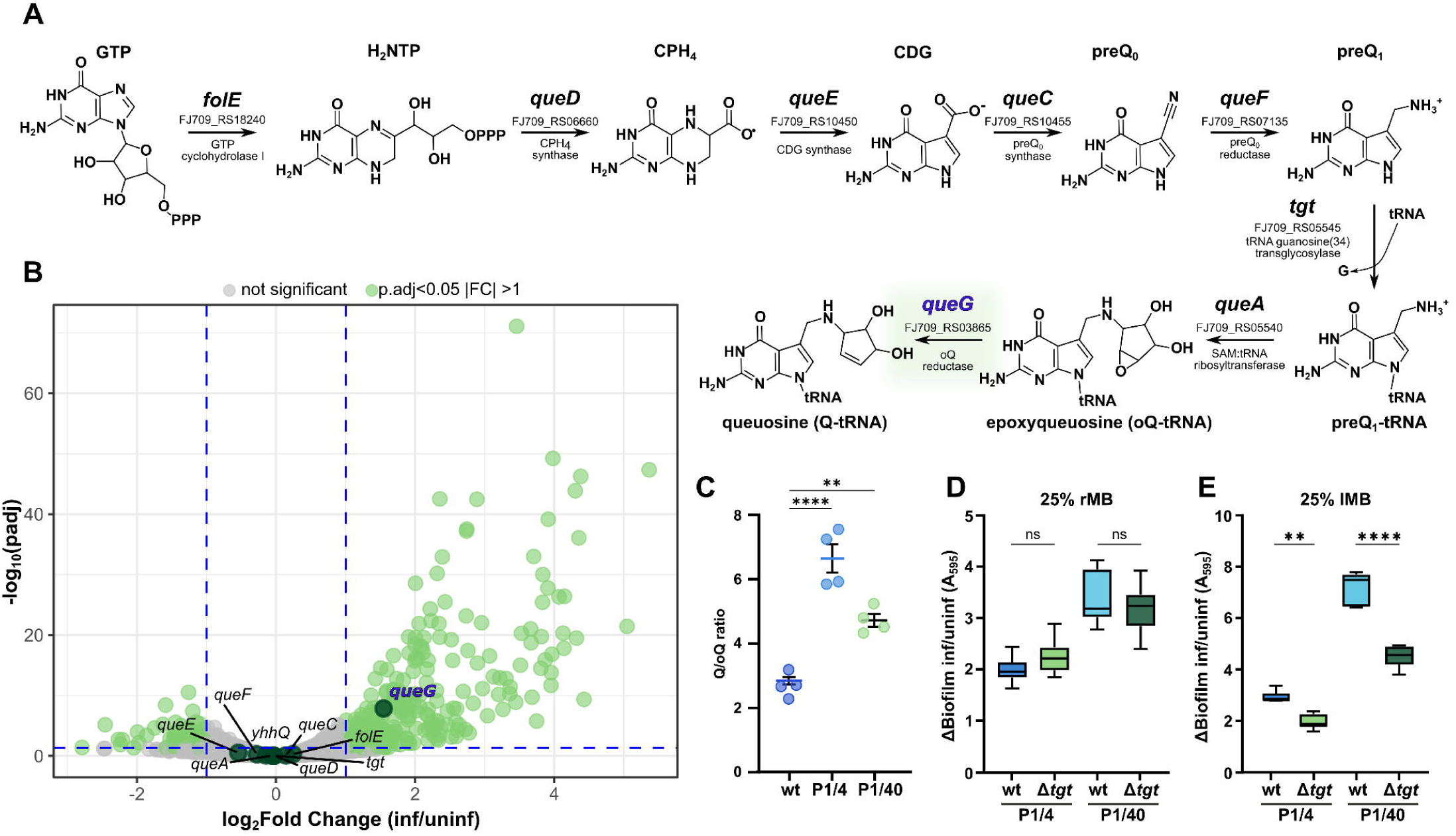
Queuosine drives higher biofilm formation during phage infection. **a**. Queuosine biosynthesis pathway in *S. glacialimarina* TZS-4_T_ with gene locus tags. Adapted from^90^. b. Volcano plot of differentially expressed genes (infected vs uninfected). Queuosine pathway genes highlighted (dark green). **c.** Quantification of queuosine (Q) and epoxyqueuosine (oQ) in tRNA by LC-MS in infected samples and uninfected control. The change of Q and oQ is expressed as ratio (Q/oQ). Ordinary one-way ANOVA with Dunnett’s multiple-comparisons correction (n = 4, ** p = 0. 0023, **** p *<* 0.0001). **d.** Difference in biofilm formation during phage infection in *S. glacialimarina* wild-type vs Δ*tgt* mutant in nutrient rich conditions (25% rMB), relative to uninfected controls (ΔBiofilm). Strains were compared using ordinary one-way ANOVA with Šidák’s multiple-comparisons correction (n = 8, ns p > 0.05). **e.** Difference in biofilm during phage infection in *S. glacialimarina* wild-type vs Δ*tgt* mutant in nutrient limited conditions (25% lMB), relative to uninfected controls (ΔBiofilm). Strains were compared using ordinary one-way ANOVA with Šidák’s multiple-comparisons correction (n = 6, ** p < 0. 01, **** p *<* 0.0001).

With this in mind, we hypothesized that the loss of Q could lead to impaired biofilm formation. To address this, we decided to create a *S. glacialimarina* deletion strain devoid of Q synthesis. Rather than attempting a *queG* deletion, which would accumulate the epoxyqueuosine (oQ) precursor, we constructed a Δ*tgt* strain that eliminates both Q– and oQ-tRNA to avoid confounding precursor effects. Initially, we compared Δ*tgt* to the wild-type strain in nutrient rich media and observed no differences in phage-induced biofilm (Fig. 2d; Suppl. Fig. 3c). Since phenotypic effects of some tRNA modifications can be masked by cellular compensation under favorable conditions^44–46^, we evaluated low-nutrient conditions to unmask potential Q-dependent phenotypes. To exclude nutrient stress effects, we first confirmed that host growth and phage replication remained comparable between the strains (Suppl. Fig. 3d-e), after which we measured the phage-induced biofilm production. Under low-nutrient conditions, Δ*tgt* showed significantly impaired biofilm induction compared to the wild-type (Fig. 2e; Suppl. Fig. 3f-g) while viable cell counts remained equivalent (Suppl. Fig. 3g), indicating that Q specifically promotes biofilm formation.

To determine whether the observed Q-dependent phenotype stems from the cumulative bias of all NAT codons or from specific ones, we subsequently examined the individual Q-decoded codons. Since oQ and Q modifications are indistinguishable on N-acryloyl-3-aminophenylboronic acid (APB) blots due to shared diol groups (Fig. 2a; Suppl. Fig. 3h), we performed functional enrichment analysis for each Q-decoded codon separately. Out of all four NAT codons, only His_CAT_ and Asp_GAT_ genes showed functional enrichment (Suppl. Fig. 3i), where His_CAT_-biased genes were enriched for biofilm-related terms and Asp_GAT_ for DNA binding and recombination functions. These results suggest that in addition to biofilm synthesis, infection-induced Q may also influence host mutagenesis.

### Mutation of *wecB* provides protection from phage infection

Given the enrichment of Asp_GAT_-biased genes in functions associated with recombination, we investigated whether Q modification influences phage resistance development. To examine the underlying mechanism, we evolved a *S. glacialimarina* strain resistant to phage 1/4 and 1/40 (Fig. 3a). While we were unable to obtain a resistant strain under continuous phage 1/40 exposure, evolution in the presence of phage 1/4 yielded a strain resistant to both phages (Suppl. Fig. 4a-b). The phage-resistant isolate displayed a pronounced granulating/flocculating phenotype, indicative of increased biofilm formation compared to the wild-type (Suppl. Fig. 4c). Collectively, these observations suggest that the resistance phenotype likely resulted from mutations in genes encoding for surface structures, which is a common mechanism underlying phage resistance^47,48^.

**Figure 3.**
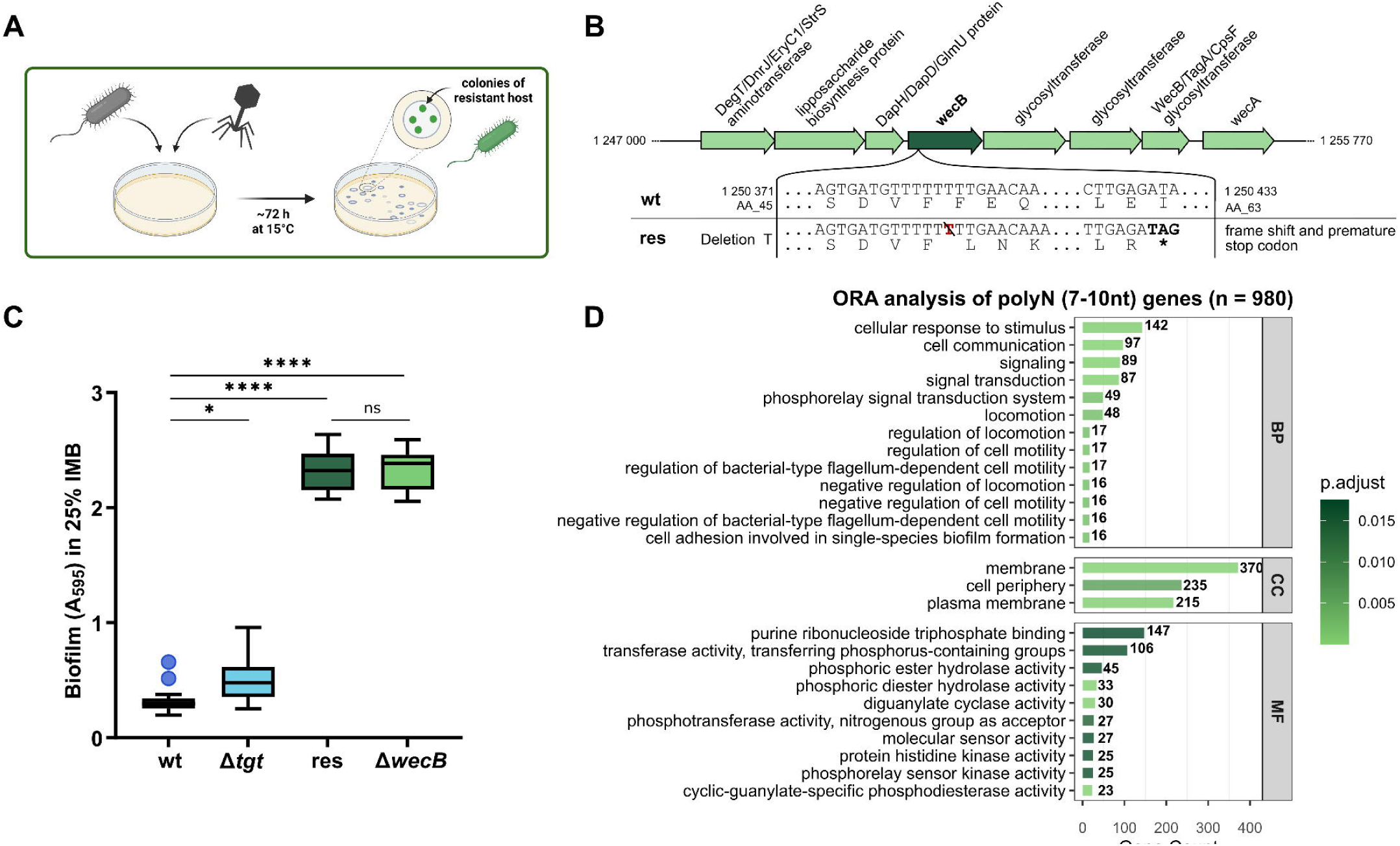
PolyN repeats in coding genes are the primary sites of host mutagenesis leading to biofilm-mediated phage resistance. **a**. Schematic representation of phage-resistant host evolution. Resistant clones were isolated from phage-cleared plaques after ∼72 h growth and further purified. Scheme created with biorender.com. **b.** Schematic of frameshift mutation in *S. glacialimarina* TZS-4_T_ phage-resistant strain (res). T-deletion in *wecB* causes frameshift and premature stop codon, yielding truncated protein (63 aa vs 357 aa full length). **c.** Biofilm production in *S. glacialimarina* TZS-4_T_ wild-type (wt), Δ*tgt,* phage-resistant (res), and Δ*wecB* strains in 25% lMB. Ordinary one-way ANOVA with Šidák’s multiple-comparisons correction (n = 16, * p < 0. 05, **** p *<* 0.0001, ns p > 0.05). **d.** GO over-representation analysis of genes containing 7-10 nucleotides long polyN stretches in coding region. Top significant GO terms (adjusted p < 0.05, Benjamini-Hochberg correction) for Biological Process (BP), Cellular Component (CC), and Molecular Function (MF) ontologies. Terms ranked by adjusted p-value.

To pinpoint the gene(s) responsible for phage resistance, we performed whole-genome sequencing of the phage-resistant strain and identified two protein-coding mutations (Supplementary Table 3): a missense substitution in a patatin-like phospholipase and a single nucleotide deletion in *wecB*, resulting in a frameshift and premature stop codon (Fig. 3b). *wecB* encodes for an epimerase that converts UDP-N-acetylglucosamine to UPD-N-acetyl-D-mannosamine as part of the enterobacterial common antigen (ECA) biosynthesis pathway^49,50^, although this pathway appears incomplete in *S. glacialimarina* (Suppl. Fig. 4d).

To ascertain whether *wecB* contributes to the observed biofilm phenotype, we generated a Δ*wecB* mutant, which exhibited increased biofilm production comparable to the resistant strain (Fig. 3c). The *wecB* mutation occurred within a polyT tract (Supplementary Table 3), suggesting a slipped-strand mispairing during DNA replication or repair. Genome-wide analysis of polyN tracts and tandem repeats revealed an enrichment of mutations in genes associated with cell surface structures, indicating these loci as potential mutational hotspots (Fig. 3d; Suppl. Fig. 4e–f; Supplementary Table 4).

### Q deficiency impairs phage resistance development

Building on our findings, we further investigated the NAT/NAC codon bias of DNA polymerases in *S. glacialimarina*, focusing on error-prone TLS polymerases. During phage infection, these polymerases are activated via the SOS response and DNA damage repair pathways^31–34^. Our analysis revealed a general bias toward NAT codons (Fig. 4a). Additionally, two of these genes, *dinB* (encoding DNA polymerase IV) and *polB* (DNA polymerase II), were also overexpressed during phage infection (Fig. 4b).

**Figure 4.**
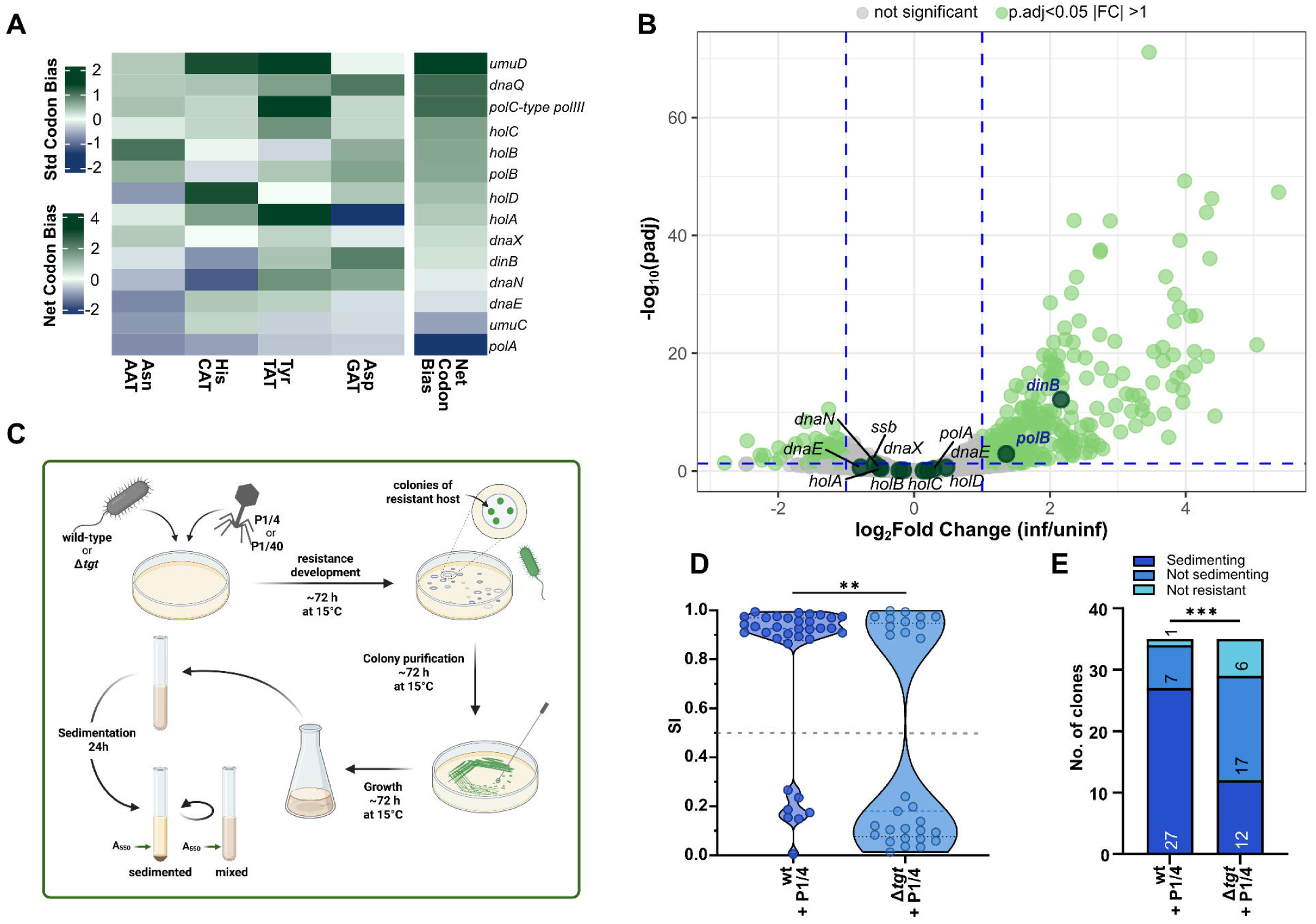
Lack of Q decreases the probability of phage resistance development via *wecB* in *S. glacialimarina* due to codon bias in DNA polymerases. **a**. Heatmap of standardized codon bias for each of Q decoded NAT codons in DNA and TLS polymerase genes (GO annotation). Net codon bias represents the sum of individual codon biases across all codons. **b.** Volcano plot of differentially expressed genes (infected vs uninfected). DNA polymerase genes from heatmap (A) highlighted. **c.** Schematic of phage-resistant *S. glacialimarina* TZS-4_T_ wt and Δ*tgt* selection and sedimentation analysis. Resistant clones were isolated from phage-cleared plaques following ∼72 h incubation, purified, grown to high density, and allowed to sediment statically for 24 h. Sedimentation index (SI) was calculated from absorbance measurements of sedimented versus homogenized cultures. Scheme created with biorender.com. **d.** Distribution of sedimentation index (SI) values for phage-resistant *S. glacialimarina* TZS-4_T_ clones from wt (n = 34) and Δ*tgt* (n = 29) strains evolved from 1/4 infected cells in nutrient limited medium. Difference of distribution was assessed by Kolmogorov-Smirnov test (** p = 0.0032). **e.** Phenotypic summary of candidate phage-resistant *S. glacialimarina* TZS-4_T_ clones developed from wt and Δ*tgt* (both n = 35) strains obtained after selection in nutrient limited medium (25% lMB). Statistical significance determined by Fisher’s exact test (*** p = 0.0008).

As TLS polymerases have been reported to be involved in generating slippage mutations^34^, their upregulation offers a potential explanation for the repetitive tract mutations observed in the phage-resistant host (Fig. 3b). We therefore expanded our codon bias analysis to SOS response and recombinational repair genes to further explore this mechanism of phage-resistance development. Our analysis revealed a general enrichment of NAT codons in TLS and repair genes (Suppl. Fig. 5a; Supplementary Table 5). Consistent with studies in other bacteria^31–33^, several genes involved in SOS response and DNA damage repair also showed elevated expression during infection (Suppl. Fig. 5b). Hence, we hypothesize that a phage-induced increase in Q modification levels, together with SOS-mediated overexpression of these genes, leads to their enhanced translation and thereby promote mutagenesis in slippage-prone repetitive tracts.

To verify the direct role of Q in this process, we quantified the sedimentation index (SI) as a proxy for the flocculating phenotype of phage-resistant mutants (Fig. 4c). Phage-resistant clones evolved in Δ*tgt* (Q-deficient) versus wild-type backbones exhibited lower SI values and resistance frequency (Fig. 4d–e; Suppl. Fig. 5c–d). Whole-genome sequencing of sedimentary phage-resistant clones from both backbones confirmed the presence of slippage mutations (e.g., in *wecB* or repetitive tracts) in most isolates (Fig. 4d–e; Supplementary Table 6). However, based on the sedimentation data, far fewer such clones evolved in the Δ*tgt* backbone (Fig. 4d–e; Suppl. Fig. 5c–d). In conclusion, the elevated slippage-mediated resistance in *S. glacialimarina* can be explained by the phage-induced increase in Q modification leading to translation enhancement of NAT-biased TLS polymerases.

### Phage resistance develops via a shared mechanism in related *Shewanella* strain

To validate our observations made in *S. glacialimarina* TZS-4_T_, we analyzed the closely related bacterium *Shewanella* sp. 40 *–* the original isolation host of phage 1/40 that was obtained concomitantly from the same location as *S. glacialimarina*^38^. The average nucleotide identity analysis (ANI) showed (Suppl. Fig. 6a) that *Shewanella* sp. 40 is a strain of *S. glacialimarina* (thereafter referred to as strain 40) rather than different species as previously reported^37,38^.

Comparative analysis of genome-wide relative synonymous codon usage (RSCU) values revealed similar codon usage patterns across both strains and their phages (Suppl. Fig. 6b). Next, we calculated NAT codon bias and identified top biased genes as described for *S. glacialimarina* strain TZS-4_T_ (Suppl. Fig. 6c and 7). In strain 40, NAT-biased genes were enriched in biofilm-associated functions, consistent with the patterns observed in TZS-4_T_ strain. However, the recombination-associated genes were not significantly enriched, likely due to a less complete functional annotation of the strain 40 genome.

Next, we performed RT-qPCR to assess the conservation of the phage-induced Q increase and the expression of SOS-induced TLS genes. Similarly to TZS-4_T_ strain, the results confirmed that *queG* and TLS genes (*dinB*, *polB, lexA*) are upregulated during phage infection in strain 40, albeit with a lower induction amplitude (Suppl. Fig. 6d). Likewise, biofilm formation also increased post-infection (Suppl. Fig. 6e). Further analysis of polyN tracts and repetitive regions in the genome showed comparable functional enrichment, suggesting similar conservation of mutagenesis hotspots (Suppl. Fig. 6f and 8).

Subsequently, we examined the *wec* pathway genes with the expectation of finding strong similarities between these *S. glacialimarina* strains. Interestingly, the strain 40 genome also contains *wecC* (Supplementary Table 7), which encodes for the UDP-N-acetyl-D-mannosamine dehydrogenase that catalyzes production of UDP-N-acetyl-D-mannosaminuronic acid^49,50^. The *wecC* gene is absent in TZS-4_T_ strain (Suppl. Fig. 4c), which leads to differences in surface ECA composition^49,50^ that may explain the naturally sediment-forming phenotype of strain 40. Consistent with this, phage-resistance evolution experiments revealed that the phage-resistant mutants predominantly exhibit non-sedimentary phenotypes Suppl. Fig. 6g-h). Whole-genome sequencing of phage-resistant clones identified slippage mutations in two out of three sequenced genomes: one in a *wecA* GTTATTG tandem repeat (1/3 genomes) and another in polyT tract of hypothetical protein with unknown function (2/3 genomes) (Supplementary Table 8).

Collectively, these findings support our proposed mechanism of phage resistance development in *Shewanella* via Q-enhanced TLS mutagenesis in repetitive hotspots, with phenotypic outcomes driven by variation in the *wec* pathway (Fig. 5).

**Figure 5.**
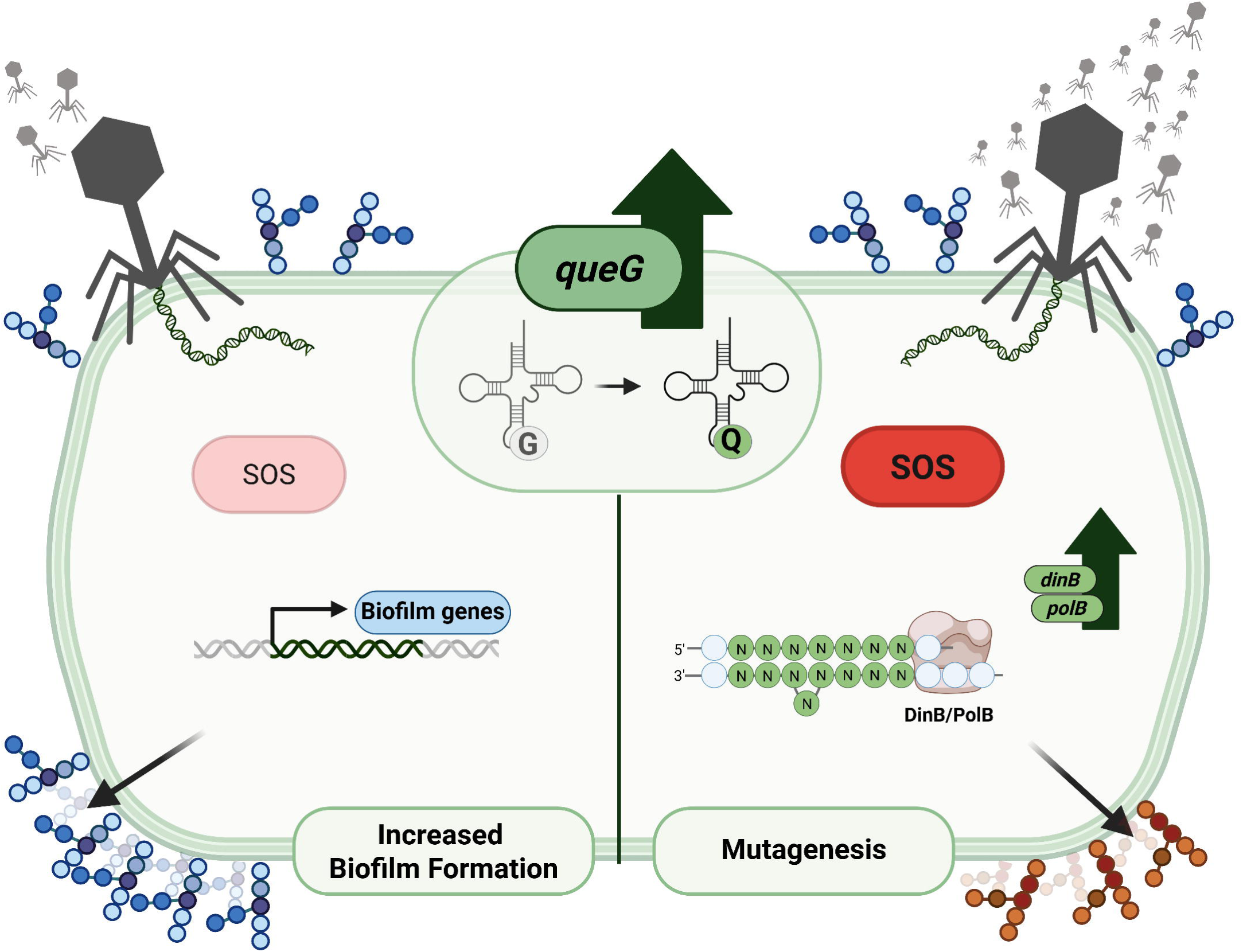
Q modification drives phage dose-dependent resistance in *S. glacialimarina*. Q-mediated enhancement of NAT codon translation promotes two phage escape routes depending on infection intensity: transient resistance via biofilm formation at low phage concentrations (phenotypic; left) and stable resistance via TLS polymerase-driven mutagenesis at high phage concentrations (genotypic; right). Scheme created with biorender.com.

## DISCUSSION

This study reveals a previously unrecognized role of phage-induced Q modification in biofilm formation and phage-resistance evolution in the marine bacterium *Shewanella glacialimarina* TZS-4_T_. Consistent with earlier findings^39^, infection with both investigated phages triggers a robust increase in Q levels, suggesting that this response constitutes a conserved host adaptation to phage stress. Furthermore, transcriptional profiling identified the epoxyqueuosine reductase *queG* as the enzyme that regulates the phage-induced increase of Q modification. Recently, expression of another Q pathway gene, *queE,* has been linked to antimicrobial peptide stress responses through increased PhoQ/PhoP signaling, although the *queE* stress-related activity appears to be unrelated to its role in RNA modification^51^. In contrast to this well-characterized regulation of *queE*, the regulatory mechanism that drives *queG* expression during phage infection remains to be investigated.

Although a functional connection between Q modification and biofilm formation has been proposed previously^13^, our data provides further mechanistic evidence linking these processes under the pressure of phage infection. While phage infection universally induces biofilm formation regardless of Q status, Q-proficient wild-type *S. glacialimarina* exhibits a significantly greater increase in biofilm production than the Q-deficient Δ*tgt* knockout strain. Additionally, our analyses revealed that the biofilm-associated genes are strongly enriched in Q-decoded NAT codons in *S. glacialimarina*. Since Q modification has been shown to enhance the translation efficiency of NAT codons^40,52^, we propose that the phage-induced increase in Q modification selectively amplifies translation of these NAT-biased genes. This mechanism reinforces biofilm matrix synthesis and reveals a previously unrecognized pathway through which phages indirectly reshape the architecture of the bacterial community.

Previous research on *Vibrio* phages indicate that phage-host interactions can lead to two distinct biofilm outcomes: a transient induction, providing temporary protection from phages, and a stable biofilm phenotype that likely arises from genetic mutations conferring phage resistance^30^. Our results mirror this duality while revealing infection dynamics as a key modulator. Specifically, low MOI-values of the more infective phage 1/4 promoted biofilm formation, whereas higher doses triggered mutagenesis and selection of stable phage-resistant variants. In contrast, for the less infective phage 1/40, even under high phage pressure we observed only biofilm development without detectable mutagenesis. This pattern aligns with recent evidence that low-level phage predation generates exogenous “danger signals”, which activate biofilm-mediated defenses in diverse bacteria without inducing mutagenesis^53^. The low infection efficiency of phage 1/40 most likely mimics such low-level predation, preferring a non-mutagenic, phenotypic strategy over genetic diversification. This illustrates how phage infection dynamics govern the balance between short-term transient phenotypic protection via biofilm formation and long-term adaptation via genetic mutation.

Beyond its role in biofilm formation, we found that Q modification appears to intersect with host mutagenic pathways through translation modulation of NAT-biased translesion synthesis (TLS) polymerases. These polymerases, typically activated by the SOS stress response, are known mediators of mutagenesis^34,35^. Phage infection has been reported to induce SOS signaling and promote TLS polymerase expression^31–33^. In *S. glacialimarina*, Q enhanced translation of NAT-biased TLS polymerases facilitated strand slippage within homopolymeric and tandem repeat-rich regions of surface-related genes, i.e. genomic sites previously identified as mutational hotspots^54,55^. These mutations generated phenotypic variants exhibiting both enhanced biofilm accumulation and phage resistance.

Among the recurrent targets of these mutations, *wecB* emerged as a critical gene. As part of the enterobacterial common antigen (ECA) biosynthesis pathway^49,50^, *wecB* contributes to surface architecture and its enzymatic activity has been reported crucial for bacteriophage N4 attachment to *E. coli*^56^. However, *S. glacialimarina* encodes only an incomplete ECA pathway and phage-induced *wecB* disruption likely causes accumulation of the *wecA*-derived intermediate undecaprenyl-phosphate *N*-acetylglucosamine (Und-P-GlcNAc). This buildup is likely to further disrupt polymer assembly and alter outer-membrane composition. In contrast, related *S. glacialimarina* strain 40 maintains a complete ECA-I lipid–like structure, and its phage-resistant mutants exhibit surface properties distinct from those of type strain *S. glacialimarina* TZS-4_T_. Taken together, these observations suggest that the *wec* pathway mediates host-phage interaction by either creating structural sites for phage attachment or, when disrupted, yielding aberrant surface polymers that promote cell aggregation and sterically hinder phage attachment. Detailed biochemical characterization of these intermediates will be essential to determine whether they act as genuine phage receptors.

Collectively, our findings present two distinct phage escape trajectories in *S. glacialimarina* (Fig. 5). A phenotypic route, driven by biofilm formation providing transient resistance, and a genotypic route yielding stable resistance. Critically, we find that Q modification orchestrates both pathways: enhancing biofilm production via enhanced translation of biofilm matrix genes, while simultaneously promoting translation of TLS polymerases enabling host genome mutagenesis. Thanks to the conservation of Q-mediated NAT codon decoding, SOS-TLS pathways and phage stress responses across bacteria, our findings establish a generalizable framework in which Q modification acts as a regulator through which bacteriophages shape bacterial adaptation.

## MATERIALS AND METHODS

### Bacterial growth conditions and phage propagation

The bacteria *Shewanella glacialimarina* TZS-4_T_ (RefSeq NZ_CM151298.1, DSM 115441)^36^ and *S. glacialimarina* 40 (RefSeq NZ_CM151295.1) were grown at +15 °C and 200 rpm in liquid media containing 25% (w/v) rich Marine broth^36^ (25% rMB; containing 7.5 g peptone (Sigma-Aldrich), 1.5 g yeast extract (Fisher BioReagents) and 9.35 g Marine broth (BD-Difco) in 1L of ddH_2_O). For limited nutrient (10× less nutrients) assays, 25% limited Marine broth (25% lMB) was used at +15 °C and 200 rpm. For complete composition of 25% lMB see Supplementary Table 9. Solid and semi-solid media were prepared by addition of 15 g/L and 7.5 g/L of agar (Fisher BioReagents), respectively.

The bacteriophage *Shewanella* sp. phage 1/4 (RefSeq NC_025436.1) and phage 1/40 (RefSeq NC_025470.1) were previously isolated from the same Baltic Sea ice sample as the host bacteria^38^. The phages were propagated by standard plate lysate method^39^ and the phage stock titer was determined by small drop plaque assay.

### Purification of Shewanella phages

Shewanella phages 1/4 and 1/40 were purified from 2 L of cleared cell lysates. To obtain cleared cell lysate, the corresponding host cells (*S. glacialimarina* TZS-4_T_ or *S. glacialimarina* 40, respectively) were grown in 25% rMB to exponential growth (OD_600_ ∼0.6) and infected with multiplicity of infection (MOI) 10 and grown until the OD_600_ declined below 0.3-0.4. The phages were precipitated from the lysate using 10% (w/v) polyethylene glycol (PEG; 6000 MW, Acros Organics) in the presence of 0.5 M NaCl (Fisher Chemical) at +4 °C for 45 min and collected by centrifugation at 8 300×*g* at +4 °C for 40 min.

Subsequently, the phage pellets were rinsed with ddH_2_O and dissolved in SM-buffer (50 mM Tris pH 7.5, 100 mM NaCl, 8 M MgSO_4_, 0.01% gelatine; all Fisher Chemical) in 1/200 volume of the original lysate, overnight at +4 °C. Next day, the viral aggregates were removed by centrifugation at 17 000×*g* for 15 min at +10 °C. The non-aggregated phages were subjected to linear 10-30% rate zonal ultracentrifugation on a sucrose gradient (in SM-buffer) at 103 700×*g* for 25 min at +10 °C. The light-scattering zone was collected and pelleted at 113 700×*g* for 3 h at 10 °C. The pelleted phages were resuspended in 0.5 mL SM-buffer and stored aliquoted at –80°C.

### Biofilm quantification

To quantify biofilm formation, the overnight cultures were diluted to OD_600_ ∼0.2 in 25% rMB or to OD_600_ ∼0.1 in 25% lMB, plated to 96-well plate and grown statically at +15 °C for 24 or 72 h, respectively. At each given incubation time point, the supernatants were removed, and cells were washed once with 200 μL of 1× PBS. Cells were fixed on plates with 200 μL 100% methanol (Fisher Chemical) for 15 min. After methanol removal, the plates were dried and stained with 200 μL of 0.05% crystal violet (Merck) for 20 min. Stain was removed and wells were washed twice with 200 μL ddH_2_O. After drying, the crystal violet was solubilized by 150 μL of 33% acetic acid (Fisher Chemical) and 100 μL of 2, 5 or 10× dilutions were prepared in ddH_2_O. The quantity of biofilm was measured as absorbance at 595 nm using Multiscan FC (ThermoFisher Scientific) plate reader.

In case of infection induced biofilm measurement, the cultures were infected with varying MOI values (0.01 for phage 1/4 and 10 for phage 1/40). In case of *S. glacialimarina* 40 infection, cells were grown to OD_600_ ∼0.2 and infected with phage 1/40 using MOI 0.01. After incubation, plates were processed as described above.

### Phage and infection characterization

#### Growth curves

Growth curves were performed by diluting 68-72 h starter cultures to OD_600_ 0.05 using 25% lMB. Then, cells were grown at +15 °C, 200 rpm and OD_600_ was monitored every 30-60 min.

#### Phage small drop plaque assay

The number of phages was determined on either 25% rMB or 25% lMB semi-solid media overlays. 250 µL or 500 μL of overnight bacterial culture was mixed with 8 mL of semi-solid agar tempered to +45 °C and poured on 12×12 cm square plate containing solid agar. Plates were dried for 20-30 min. For each phage sample, 10-fold serial dilutions in SM-buffer were prepared and 10 μL drops were put on the prepared bacterial layer. After the drops dried, plates were incubated overnight at +15 °C. The number of phages was determined by counting plaques and calculating number of plaques per 1 mL (PFU/mL).

#### Viable cells counting

To determine the viable count (VC), the cultures were sampled at different times throughout growth. Culture OD_600_ was measured, culture sample was serially diluted, plated as 10 μL drops on 12×12 cm square plates containing 25% rMB or 25% lMB agar and plates were grown for 48 h or 72 h, respectively. VC was calculated as number of colony forming units per 1 mL (CFU/mL).

#### 1-step growth curve

One-step growth curves were performed as described^57^ with few modifications. Host cultures were grown to OD_600_ 0.6 or 0.3 (rMB or lMB), mixed with 10^6^ PFU of phage and incubated for 5 min at +15 °C. After incubation, the adsorption control was prepared by collecting 100 μL of culture and mixing it with 5 μL chloroform. The remaining culture was serially 10-fold diluted into new flasks and incubated at +15 °C, 200 rpm. The culture was sampled by withdrawing 100 μL every 10-60 min and phages were detected by direct plating of 10 μL drops on 12×12 cm square plates as described above.

#### Phage adsorption assay

Exponentially growing cultures (OD_600_ ∼0.6 for 25% rMB or ∼0.3 for 25% lMB) were infected by ∼ 300 000 phages, whereas control flasks contained only medium and phages. The culture and control flasks were left shaking at 70 rpm and sampled at 0-45 min. 100 μL samples were collected into tubes with 850 μL 25% rMB and 50 μL chloroform. Alternatively, 100 μL samples were collected into tubes with 900 μL and bacteria was removed by centrifugation at 13 000×*g*, 1 min, +4 °C. The number of free phages was determined by small drop plaque assay as described above and adsorption was calculated as % of adsorbed phages.

### Gene knockout preparation

#### Vector preparation

The gene knockouts were prepared using SIBR system^58^ cloned to a new *S. glacialimarina* compatible minimal expression vector pFSD.

First, the pFSD was prepared by removal of conjugation elements and insertion of MCS from of broad host range vector pJRD215^59^, a gift from Renato Morona (Addgene plasmid #167555; http://n2t.net/addgene:167555). The desired elements (OriV, Rep genes, Neo/Kan resistance genes, promoter region of rep genes) were amplified by PCR with 20 bp overlaps. The purified PCR fragments were ligated together using TEDA assembly [100 mM Tris-HCl pH 7.5, 10 mM MgCl_2_, 10 mM dithiothreitol, 5% PEG-8000, 0.002 U/μL T5 exonuclease (NEB)] as previously described^60^. The resulting plasmid was further mutated to remove remaining mobilizable elements using site-directed mutagenesis. The MCS was amplified by PCR from pX1^61^, a gift from Monica Hollstein (Addgene plasmid #46848; http://n2t.net/addgene:46848) plasmid and inserted to pFSD via restriction-insertion using *Smi*I. Final plasmid was sequenced by whole plasmid sequencing (Eurofins). The pFSD plasmid is available from Addgene (#250975).

Next, the pFSD was further modified by inserting SIBR cassette via *Mun*I from plasmid SIBR0004-NTBsaI which was generated by spacer exchange from SIBR0004^58^, a gift from John van der Oost (Addgene plasmid #177667; http://n2t.net/addgene:177667). The resulting plasmid pFSD-SIBR0004 was sequenced (Eurofins) and served as empty vector for introducing homologous regions and spacer for counterselection of successful mutants. pFSD-SIBR004 is available from Addgene (#250974).

Plasmids for Δ*tgt* and Δ*wecB* knockouts were generated by two step cloning. First, each respective spacer sequence was inserted to pFSD-SIBR0004 via *Bsa*I. The spacer inserts were prepared by annealing two oligonucleotides with *Bsa*I compatible overhangs. Second, the upstream and downstream 700-800 bp homologous regions of gene *tgt* (locus tag FJ709_RS05545) or *wecB* (FJ709_RS05430) were PCR amplified, purified and ligated to *Eco*RI/*Bam*HI linearized pFSD-SIBR004 containing spacer using TEDA assembly. The plasmids (pFSD-SIBR0004-tgt and pFSD-SIBR0004-wecB) were sequence verified by whole plasmid sequencing (Eurofins).

All colony PCRs were performed using 5x Hot FirePol mix (Solis BioDyne) according to manufacturer’s instructions. Primers used for cloning and spacers are in Supplementary Table 10.

#### Preparation of electrocompetent cells

To generate knockouts, the pFSD-SIBR0004-tgt/wecB plasmids were introduced to *S. glacialimarina* TZS-4_T_ by electroporation. The electroporation cells were prepared as follows. Starter cultures of *S. glacialimarina* grew in 20 mL 25% rMB at +15 °C, 200 rpm. The 34-36 h culture was used to inoculate 100 mL of 25% rMB to OD_600_ ∼0.2 and grown at +15 °C shaking at 200 rpm until an OD_600_ of 0.3-0.4. Then, the culture was cooled down on ice and cells were harvested by centrifugation at 3 000×*g*, 10 min at +4 °C. Supernatant was removed and cells were resuspended in 50 mL ice-cold wash buffer (10 mM MgCl_2_, 5 mM CaCl_2_) followed by centrifugation at 3 000×*g*, 10 min at +4 °C. The wash step was repeated once more, and the final cell pellet was resuspended in 1 mL of ice-cold 1M sorbitol (Fisher Chemical). The cell suspension was aliquoted (50 μL), flash-frozen in liquid nitrogen and stored at –80 °C prior to transformation.

#### Vector transformation and knockout screening

Knockout vectors (2 μg) were electroporated to 50 μL *S. glacialimarina* competent cells in pre-cooled 0.1 cm cuvettes using Gene Pulser (Biorad) using 12.5 kV/cm, 800 Ω. Cells were recovered in 1 mL of 25% rMB for 2 h in table-top thermomixer (Eppendorf) at +15 °C and 900 rpm. Due to low electroporation and genome editing efficiency, recovered cells were transferred to 9 mL 25% rMB with 50 μg/mL kanamycin and grown for 24-72h. Every 24h post electroporation, 100 μL of culture was serial diluted and spotted on 25% rMB plates containing 50 μg/mL kanamycin and 1 mM theophylline to activate SIBR system for successful knockout selection. Plates were grown for 48 h at +15 °C. The colonies were tested by colony PCR using Phusion polymerase mix (ThermoFisher Scientific) and primers upstream and downstream of *tgt* or *wecB* gene, respectively. Primers used for colony screening are in Supplementary Table 10. Positive clones were grown in 25% rMB and mutation was verified by whole genome sequencing (Eurofins).

### RT-qPCR

Gene expression was evaluated by RT-qPCR using Pfaffl ddCt method^62^. Briefly, total RNA from infected and uninfected *Shewanella glacialimarina* TZS-4_T_ and 40, was DNase treated, and 2 μg were reverse transcribed by MaximaRT (ThermoFisher Scientific) using random hexamers according to manufacturer’s instructions. cDNA was diluted 50× and 1 μL was used per 10 μL reaction containing 500 nM of each primer and 1x PerfeCTa SYBR Green FastMix (Quantabio). All RT-qPCR runs were performed on Quantstudio3 (ThermoFisher Scientific). The list of tested genes and used primers is in Supplementary Table 10.

### Development of phage-resistant bacterial hosts

To develop phage resistant *S. glacialimarina* TZS-4_T_ (wt and Δ*tgt)* and *S. glacialimarina* 40 strains, the host lawns were prepared on 25% rMB or lMB semi-solid agar as described above. After plates solidified, 20 μL drops containing high amount of purified phage 1/4 (∼10^9^ phages) were spotted on the host and let to dry. After the droplets dried, the plate was incubated at +15 °C for 3 days until first resistant colonies appeared. Single colonies were picked and re-stroke on fresh 25% rMB/lMB agar plate to further purify the clones. The single colony from each clone was then propagated in 25% rMB/lMB and stored as glycerol stocks at –80 °C. The resistance was verified by spotting phages on resistant host lawn as described above.

To analyze mutations, genomic DNA was isolated from overnight cultures using Monarch^®^ Spin gDNA Extraction Kit (NEB) and sequenced by ONT long read sequencing (Eurofins). The genomes were assembled using Flye^63^ (v2.9.3) with parameters optimized for bacterial genomes. Genomes were aligned to reference genomes of *S. glacialimarina* TZS-4_T_ or 40 genome using Nucdiff^64^ (v2.0.3).

### Measurement of sedimentation index

To measure sedimentation of new mutant clones, we propagated each clone in 20 mL of 25% rMB (strain 40) or 25% lMB (all other strains) for 34-36 or 70-72 h at +15 °C, 200 rpm to obtain dense cultures. 5 mL of each grown culture was transferred to glass test tubes, mixed thoroughly, and let to sediment at room temperature for 24 h. The sedimentation was measured as A_550_ (sedimented) using Clormic (J.P.Selecta), tubes were then vortexed and measured again (mixed) to control for variability of growth. The sedimentation index (SI) was calculated as (A_550_^mixed^-A_550_^sedimented^)/A_550_^mixed^. The clones which SI > 0.5 were considered to have sedimenting phenotype, while those with SI < 0.5 were not sedimenting. All SI values are in Supplementary Table 11.

Genomic DNA isolation and sequencing of phage resistant *S. glacialimarina* TZS-4_T_ **and strain 40**

Genomic DNA was isolated from overnight cultures of *S. glacialimarina* TZS-4_T_ phage resistant mutant (res) and 40 strains using Monarch^®^ Spin gDNA Extraction Kit (NEB) and sequenced by PacBio Revio using HiFi long read sequencing (DNA Sequencing and Genomics Laboratory, Institute of Biotechnology, University of Helsinki). Genomes were assembled by Flye assembler^63^ (v2.9.5-b1801) using 80 000 reads HiFi reads per genome. Assembled genomes were annotated with Bakta^65^ (v1.11). The completeness of the genomes was checked by performing BUSCO^66^ (v5.8.3). *S. glacialimarina* 40 was compared to *S. glacialimarina* TZS-4_T_ using fastANI^67^ (v.1.34). Map with comparison track was created with Proksee^68^.

### RNA and tRNA isolation

#### Total RNA isolation from phage infected cells

Total RNA was isolated as described before^39^ with few modifications. For RNA isolation, 20 mL cultures were infected with MOI 10 (infected) or SM-buffer (control), and cells were harvested at 70 min post infection (p.i.). The cell pellet was resuspended in 8 mL of Trizol (0.8M guanidine thiocyanate, 0.4M ammonium thiocyanate, 38% acidic phenol, 0.1 M sodium acetate, 5% glycerol) with addition of 0.8 mL of 3-bromo-1-chloropropane (BCP, Acros Organics) and glass beads. The mixture was vortexed for ∼5 min and centrifuged at 10 000×*g*, 15 min, 22 °C. The RNA was further re-isolated from the aqueous phase by another round of 4 mL Trizol and 0.4 mL BCP. The aqueous phase was finally re-extracted by addition of 2 mL acidic phenol (pH 4, Sigma-Aldrich), 400 µL BCP. Then, RNA was precipitated from the aqueous phase with 99.6% ethanol. The RNA pellets were air-dried and dissolved in ddH_2_O.

#### tRNA isolation

tRNA was isolated by silica spin columns as described^69^ using 400 μg of total RNA. The final tRNA was eluted in ddH_2_O and stored at –80 °C. The tRNA purity was assessed by electrophoresis on denaturing 10% 8 M urea-polyacrylamide gels.

### Queuosine detection and quantification

#### Q quantification by UHPLC-MS

Dephosphorylated monoribonucleosides (1000 ng) were prepared and analyzed by UHPLC-MS as previously described^70^ using a SYNAPT G2-Si HRMS system (Waters). MS data analysis was carried out in MZmine^71^, v4.8. Queuosine (Q) and epoxyqueuosine (oQ) were identified based on ion masses reported in Modomics^72^. The quantities of Q and oQ are expressed as peak area normalized to m^1,3^Ψ spiking. Normalized values for Q and oQ are in Supplementary Table 12.

#### Q detection by Northern blot

Detection of Q was performed by chemiluminescent Northern blot using APB gels as previously described^69^. The sequences of used probes are listed in Supplementary Table 13.

### RNA-seq library preparation and data analysis

#### RNA-seq library preparation

Total RNA from 70 min post infection was isolated as described above, DNase-treated, reisolated with two rounds of acidic phenol (pH 4) and BCP. Then, the total RNA was rRNA depleted using RiboCop rRNA depletion kit for G-bacteria (Lexogen) and RNA-seq libraries were prepared by Corall RNA-seq V2 with UDI 12 nt (Lexogen) according to manufacturer’s protocol. Final libraries were quantified, pooled and sequenced on Illumina NovaSeq6000 to generate 50 bp paired-end reads (Next Generation Sequencing Facility at Vienna BioCenter Core Facilities).

#### RNA-seq data analysis

Data were demultiplexed (Vienna Biocenter) and quality was checked by fastQC^73^ (v0.12.1). UMIs were extracted using regex with UMItools^74^ (v1.1.5). Reads were then trimmed for polyG, adapters and quality trimmed with Cutadapt^75^ (v5.1) and fastp^76^ (v0.24.0). Remaining reads were mapped to merged host and phage reference genomes with Bowtie2^77^ (v2.5.4), coordinate sorted and converted to bam with SAMtools^78^ (v1.21). Next, mapped reads were UMI deduplicated using UMItools dedup. Multimapping and improperly paired reads were filtered out using SAMtools^78^ (v1.21). Finally, read counts for each sample were called with featureCounts^79^ from subread package (v2.0.6).

Differential expression analysis was done in R (v4.5.2) using custom scripts. DESeq analysis was performed with DESeq2 package^80^ (v1.50.2) with an adjusted p-value threshold of <0.05. Viral reads were removed post DESeq2 analysis. For visualization, log_2_ fold changes were shrunk using apeglm^81^ (v1.32.0). Complete DESeq2 results are in Supplementary Table 14.

### Additional data and computational analyses

#### Functional genome re-annotation of S. glacialimarina

To improve annotation, we re-sequenced *S. glacialimarina* TZS-4_T_ genome using ONT sequencing (Eurofins). Genome was assembled with Flye^63^ (v2.9.3) and annotated with Bakta^65^ (v1.11). The new locus tags were cross mapped with existing RefSeq annotations (NZ_CP041216.1 and NZ_CM151298.1). The functional annotation of predicted protein coding regions (CDS) was performed via orthology assignment with eggNOG-mapper^82^ (v2.1.13), function prediction by InterProScan^83^ (v5.76.107.0) and Pannzer2^84^. The GO and COG term annotations were combined, and non-redundant OrgDb database was created using AnnotationForge^85^ (v1.52.0) for use with clusterProfiler^86^ (v4.18.1). A similar procedure was applied for the *S. glacialimarina* 40 genome. Complete functional annotation of both hosts is in Supplementary Tables 15 and 16.

#### Analysis of codon usage

The standardized codon usage bias was calculated essentially as described previously^3^. First, the CDS sequences were extracted from genome using gffread^87^ (v0.12.7). The sequences were imported to R and cleaned for proper stop/start codons and pseudogenes removal using Cubar package^88^ (v1.2.0). The codons per gene were counted and the proportion of each codon per amino acid were calculated per genome and gene. The codon usage bias for a given codon and a gene was calculated by subtracting the codon proportion per whole genome from the codon proportion obtained for the gene. Standardized codon usage bias was calculated by dividing the difference by standard deviation of these differences across all genes. The top codon biased genes were identified per codon using kernel density estimation (KDE) with Epanechnikov kernels. Bandwidth was selected as the median of three standard methods: Sheather-Jones (bw.SJ), biased cross-validation (bw.bcv), and normal reference rule (bw.nrd0), preventing instability from individual selector failures. The threshold was the rightmost local density minimum (75^th^ percentile fallback if none detected); genes exceeding this per-codon cutoff were deemed as most biased (Supplementary Tables 17-18). The list of the top biased genes was then analyzed by over-representation analysis using clusterProfiler^86^ (v4.18.1). Net codon bias was calculated as gene-wise sum of all standardized codon biases for NAT codons (Supplementary Table 17-18). RSCU values for host and phage genomes were calculated by cubar^88^ (v1.2.0).

#### Genome-wide PolyN and tandem repeat analysis

The sequences and annotations of the host genomes were imported to R using Biostrings^89^ (v2.78.0). Using a custom R function, the number of 7-10 nt long polyN in coding regions were counted. Additionally, tandem repeats of minimum 5 nt long were identified (Supplementary Table 4 and 19). The list of genes the polyN regions and tandem repeats were analyzed by over-representation analysis using clusterProfiler^86^ (v4.18.1).

#### Statistical analyses and visualization

Statistical analyses and visualization were performed using GraphPad Prism (v10.1.2), R (v4.5.2, http://www.r-project.org) and Inkscape (v1.4.2). Parametric tests used when data met assumptions of normality (Shapiro-Wilk) and equal variances (Bartlett’s); otherwise, Brown-Forsythe ANOVA or non-parametric alternatives applied. Two tailed tests were applied throughout, unless otherwise stated. More details on specific statistical tests as well as the number of biological replicates are denoted in each figure legend.

## DATA AVAILABILITY

All sequencing data described in this study are publicly available through NCBI. The raw sequencing data from RNA-seq data were deposited on NCBI under project PRJNA1416385 and raw counts to GEO GSE329327. The genome sequences are available under project accession number PRJNA1299150. The generated Bakta annotation gff files are available on GitHub repository https://github.com/PavlinaG/ShewanellaBiofilmAndQ.

## CODE AVAILABILILITY

The used code for calculation of standardize codon bias, RNA-seq and repetitive regions analysis is available via GitHub at https://github.com/PavlinaG/ShewanellaBiofilmAndQ.

## ACKNOWLEDGEMENTS

This work was supported by the Research Council of Finland [grant no. 354906 to L.P.S.], the Novo Nordisk Foundation [grant no. NNF19OC0054454 to L.P.S.], and the Sigrid Jusélius Foundation [grant no. 240192 to L.P.S.]. P.G. is a fellow of Doctoral Programme in Integrative Life Science, University of Helsinki. The authors thank Dr. Elina Roine for providing the *Shewanella* strains and phages, as well as the HiLIFE Biocomplex Unit, University of Helsinki—a member of Instruct-ERIC Centre Finland, FINStruct, and Biocenter Finland—for high-speed and ultracentrifugation services. RNA sequencing was performed by the Next Generation Sequencing Facility at Vienna BioCenter Core Facilities (VBCF), member of the Vienna BioCenter (VBC), Austria. The authors are grateful to Miikka Olin, Department of Food and Nutrition, Faculty of Agriculture and Forestry, University of Helsinki for the use of their Synapt G2 Si mass spectrometer. Lastly, we acknowledge the DNA Sequencing and Genomics Laboratory (supported by HiLIFE and Biocenter Finland), Institute of Biotechnology, University of Helsinki for whole-genome sequencing and genome assembly.

## AUTHOR CONTRIBUTIONS

Conceptualization – P.G., L.P.S.; Data curation – P.G., M.K.H.; Formal analysis – P.G., M.K.H., L.P.S.; Funding Acquisition – L.P.S.; Investigation – P.G., M.K.H., N.S.; Methodology – P.G., M.K.H., N.S.; Project Administration – L.P.S.; Supervision: L.P.S.; Visualization – P.G., M.K.H.; Writing – original draft – P.G.; Writing – Reviewing & Editing – P.G., M.K.H., N.S., L.P.S.

## SUPPLEMENTARY INFORMATION

Supplementary Tables 1-19

Supplementary Figures 1-8

## FIGURES AND FIGURE LEGENDS

**Suppl. Figure 1.** Infection of *S. glacialimarina* TZS-4_T_ by phages 1/4 and 1/40 and codon biases of host biofilm genes. **a.** Titers of phage 1/40 on *S. glacialimarina* TZS-4_T_ and its isolation host *Shewanella* sp. 40 (left) and corresponding efficiency of plaquing (EOP) of phage 1/40 on *S. glacialimarina* (right). Colored lines and error bars represent mean and s.e.m. (n = 6), respectively. **b.** One-step growth curves of phages 1/4 and 1/40 infecting *S. glacialimarina* TZS-4_T_. Error bars represent s.e.m. (n = 3). **c.** Violin plots showing distribution of standardized codon bias for NAC and NAT codons across all coding genes. **d.** Over-representation analysis of same top biased genes for NAC (322 genes) and NAT (1302 genes) codons as in Figure 1d using COG. The significant COGs (adjusted p < 0.05, Benjamini-Hochberg correction) are ranked by adjusted p-value. **e.** Heatmap of standardized codon bias for each of Q decoded NAT codons in genes involved in biofilm formation (n = 353, GO and COG annotation). Net codon bias represents the sum of individual codon biases across all codons. List of genes is in Supplementary Table 2.

**Suppl. Figure 2.** Thresholds for selection of top NAT and NAC codon biased protein coding genes in *Shewanella glacialimarina* TZS-4_T_. Number of codon biased genes for each codon is shown above each graph, and red line indicates each threshold value. Genes on right are considered top biased. **a.** Threshold for codons coding for asparagine. **b.** Thresholds for codons coding of histidine. **c.** Thresholds for codons coding for aspartic acid. **d.** Thresholds for codons coding for tyrosine.

**Suppl. Figure 3.** Queuosine effect on biofilm formation during phage infection. **a.** Expression patterns of Q pathway genes (as in Fig. 2a) shown as log₂ fold changes (±95% CI) during phage 1/4 infection of *S. glacialimarina* TZS-4_T_ at 70 min post-infection (p.i.). Epoxyqueuosine (oQ) reductase *queG* is highlighted in dark green. **b.** Heatmap of mean normalized counts (DESeq2) ± s.e.m. for queuosine pathway genes (panel a) (n = 4). **c.** Biofilm formation during phage infection in *S. glacialimarina* wild-type and Δ*tgt* strains under nutrient-rich conditions (25% rMB), as in Fig. 2d. Biofilm levels compared by one-way ANOVA with Šidák’s multiple-comparisons correction (n = 8; ****p < 0.0001, ns p > 0.05). **d.** Growth curves of *S. glacialimarina* wild-type and Δ*tgt* strains in limited nutrient medium 25% lMB (left; mean ± s.e.m., n = 3) and cell counts vs. OD₆₀₀ (right; mean ± s.e.m., n = 3). **e.** Adsorption of phage 1/4 on host *S. glacialimarina* wild-type and Δ*tgt* strains (left) and one-step growth curve (right; mean ± s.e.m., n = 3). Adsorption was compared by unpaired t-test. **f.** Biofilm formation during phage infection in *S. glacialimarina* wild-type and Δ*tgt* strains under nutrient limited conditions (25% lMB), as in Fig. 2e. Biofilm levels compared by Brown-Forsythe ANOVA with Dunnett’s multiple-comparisons correction (n = 6; * p < 0.05, **** p < 0.0001, ns p > 0.05). **g.** Viable counts from biofilm assay (as in panel f) for cells in biofilms vs. planktonic phase in 25% lMB. Cell counts were compared by two-way ANOVA with Šidák’s multiple-comparisons correction (mean ± s.e.m., n = 3). **h.** APB Northern blot analysis of queuosine-carrying tRNA isoacceptors in *S. glacialimarina* wild-type and Δ*tgt* strains during phage infection. Q denotes signal corresponding to queuosine and G to unmodified guanosine. **i.** GO over-representation analysis of top biased genes for NAT (1302 genes) codon split by amino acid. Top significant GO terms (adjusted p < 0.05, Benjamini-Hochberg correction) for Biological Process (BP), Cellular Component (CC), and Molecular Function (MF) ontologies. Terms ranked by adjusted p-value.

**Suppl. Figure 4.** The *wecB* mutation confers phage resistance in *S. glacialimarina* TZS-4_T_. **a.** Phage 1/4 adsorption assay for wild-type and phage-resistant (res) strains of *S. glacialimarina* (left; mean ± s.e.m., n = 3). One-step growth curve of phage 1/4 on wild-type and res strains (right; mean ± s.e.m., n = 3). b. Phage 1/4 and 1/40 titration result on host *S. glacialimarina* wild type and resistant strains. **c.** Viable counts vs. OD_600_ for *S. glacialimarina* phage-resistant (res) and wild-type strains (mean ± s.e.m., 3 biological replicates). **d.** Schematic of the *wec* pathway in *S. glacialimarina*, a typical part of enterobacterial common antigen (ECA) synthesis, with only ECA Lipid I genes identified. **e.** Distribution of polyN regions in the *S. glacialimarina* genome. The exact number of instances for each category is shown in columns. **f.** GO over-representation analysis of genes (n = 2329) containing >5 nt long tandem repeats in coding region. Top significant GO terms (adjusted p < 0.05, Benjamini-Hochberg correction) for Biological Process (BP), Cellular Component (CC), and Molecular Function (MF) ontologies. Terms ranked by adjusted p-value.

**Suppl. Figure 5.** SOS response, DNA repair, and recombination genes involved in phage resistance development in *S. glacialimarina* TZS-4_T_ are biased toward NAT codons. **a**. Heatmap of standardized codon bias for each of Q decoded NAT codons in genes involved in SOS response, recombination and damage repair (n = 138) in *S. glacialimarina* (GO). Net codon bias represents the sum of individual codon biases across all codons. List of genes is in Supplementary Table 5. **b.** Volcano plot of differentially expressed genes (infected vs uninfected). SOS response associated genes from heatmap (A) are highlighted. **c.** Example picture of sedimented phage-resistant wild-type and Δ*tgt S. glacialimarina* strains used for sedimentation index (SI) calculation in Figure 4d-e. **d.** Distribution of sedimentation index (SI) values for all tested strains (n = 5 for control strains and phage 1/40 infected; n = 35 for phage 1/4 infected).

**Suppl. Figure 6.** Phage-resistance develops via NAT-biased genes in related *Shewanella glacialimarina* strain 40 by similar mechanism. **a.** BLAST comparison (outer green track) of the genomes of *S. glacialimarina* 40 (sp. 40) and *S. glacialimarina* TZS-4_T_ (wt). ANI score was calculated by fastANI^67^. Map with comparison track was created with Proksee^68^. **b.** Correlation of RSCU values between host and phage genomes. **c.** GO over-representation analysis of top biased genes for NAT (1269 genes) codon in *S. glacialimarina* 40 (sp. 40) split by amino acid. Top significant GO terms (adjusted p < 0.05, Benjamini-Hochberg correction) for Biological Process (BP) and Cellular Component (CC) ontologies. Terms ranked by adjusted p-value. **d.** RT-qPCR analysis of SOS and Q pathway genes expression during phage infection in hosts *S. glacialimarina* 40 (strain 40; light blue) and type strain (wt; dark blue). Dotted line crosses fold change = 1 (no change). **e.** Biofilm formation during phage 1/40 infection (MOI = 0.01) in *S. glacialimarina* strain 40 (strain 40) under nutrient-rich conditions (25% rMB). Biofilm levels compared by unpaired t-test (n = 8; ****p = 0.0001). **f.** Distribution of polyN regions in the *S. glacialimarina* strain 40 (strain 40) genes compared to *S. glacialimarina* TZS-4_T_ (wt). The exact number of instances for each category is shown in columns. **g.** Phenotypic summary of candidate phage-resistant *S. glacialimarina* strain 40 clones (n = 20) developed in nutrient rich medium (25% rMB). **h.** Distribution of sedimentation index (SI) values for phage-resistant strain 40 clones evolved from phage 1/40 infected cells. Difference in distributions was assessed by Kolmogorov-Smirnov test (n = 5 control, n = 14 resistant, ** p = 0.001).

**Suppl. Figure 7.** Thresholds for selection of top NAT and NAC codon biased protein coding genes in *Shewanella glacialimarina* 40. Number of codon biased genes for each codon is shown above each graph, and red line indicates each threshold value. Genes on right are considered top biased. **a.** Threshold for codons coding for asparagine. **b.** Thresholds for codons coding of histidine. **c.** Thresholds for codons coding for aspartic acid. **d.** Thresholds for codons coding for tyrosine.

**Suppl. Figure 8.** Over-representation analysis of coding genes containing polyN and tandem repeats in *Shewanella glacialimarina* 40. **a.** ORA analysis of coding genes containing 7-10 nt long polyN regions. Top significant GO terms (adjusted p < 0.05, Benjamini-Hochberg correction) for Biological Process (BP), Cellular Component (CC), and Molecular Function (MF) ontologies. Terms ranked by adjusted p-value. **b.** ORA analysis of coding genes containing >5 and <15nt long tandem repeats. Top significant GO terms (adjusted p < 0.05, Benjamini-Hochberg correction) for Biological Process (BP), Cellular Component (CC), and Molecular Function (MF) ontologies. Terms ranked by adjusted p-value.

