## Supplementary figures and images for "Queuosine promotes *wecB*-dependent phage resistance and biofilm formation in the marine bacterium *Shewanella glacialimarina*"

### Supplementary Figure 1

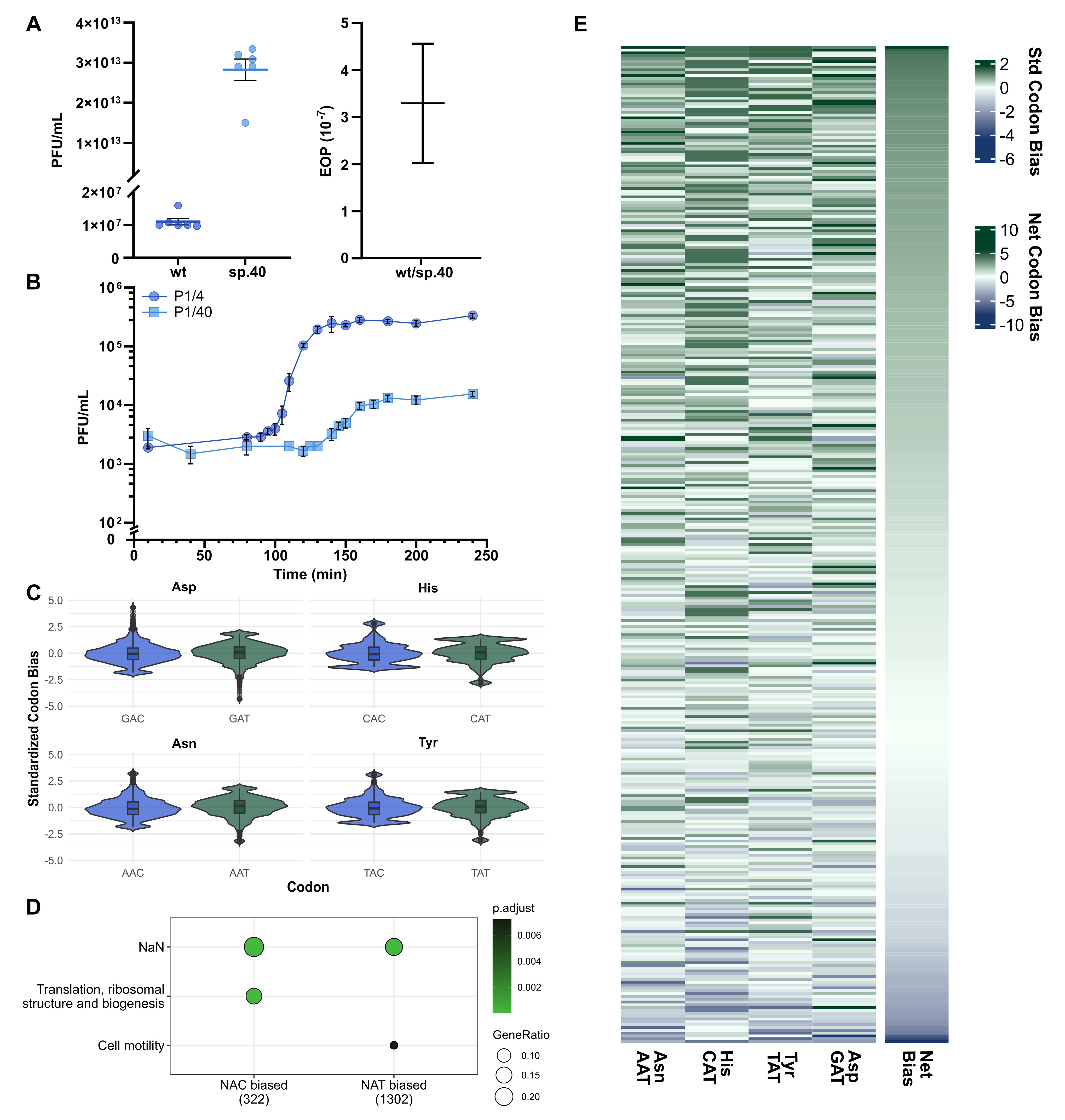

### Supplementary Figure 2

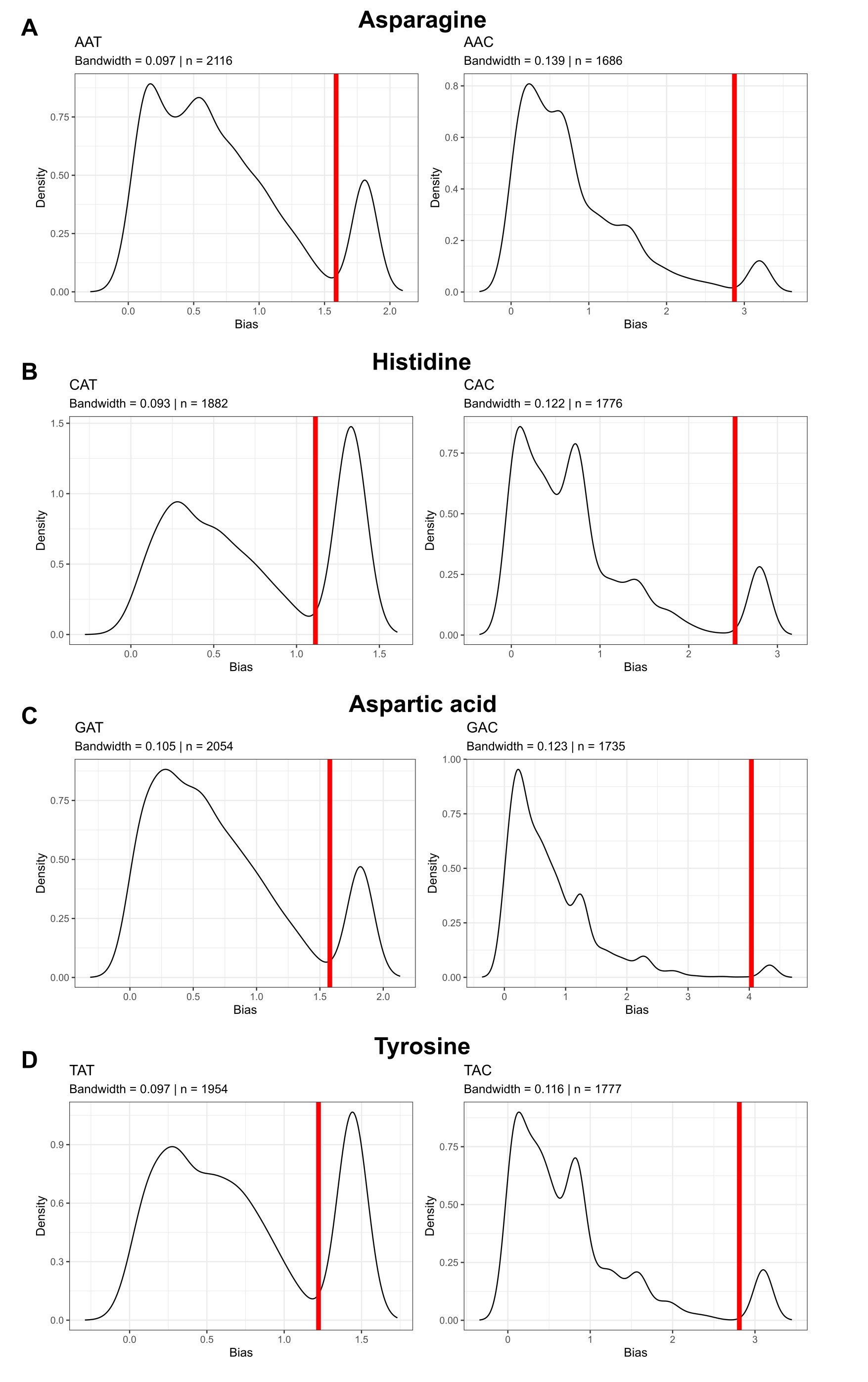

### Supplementary Figure 3

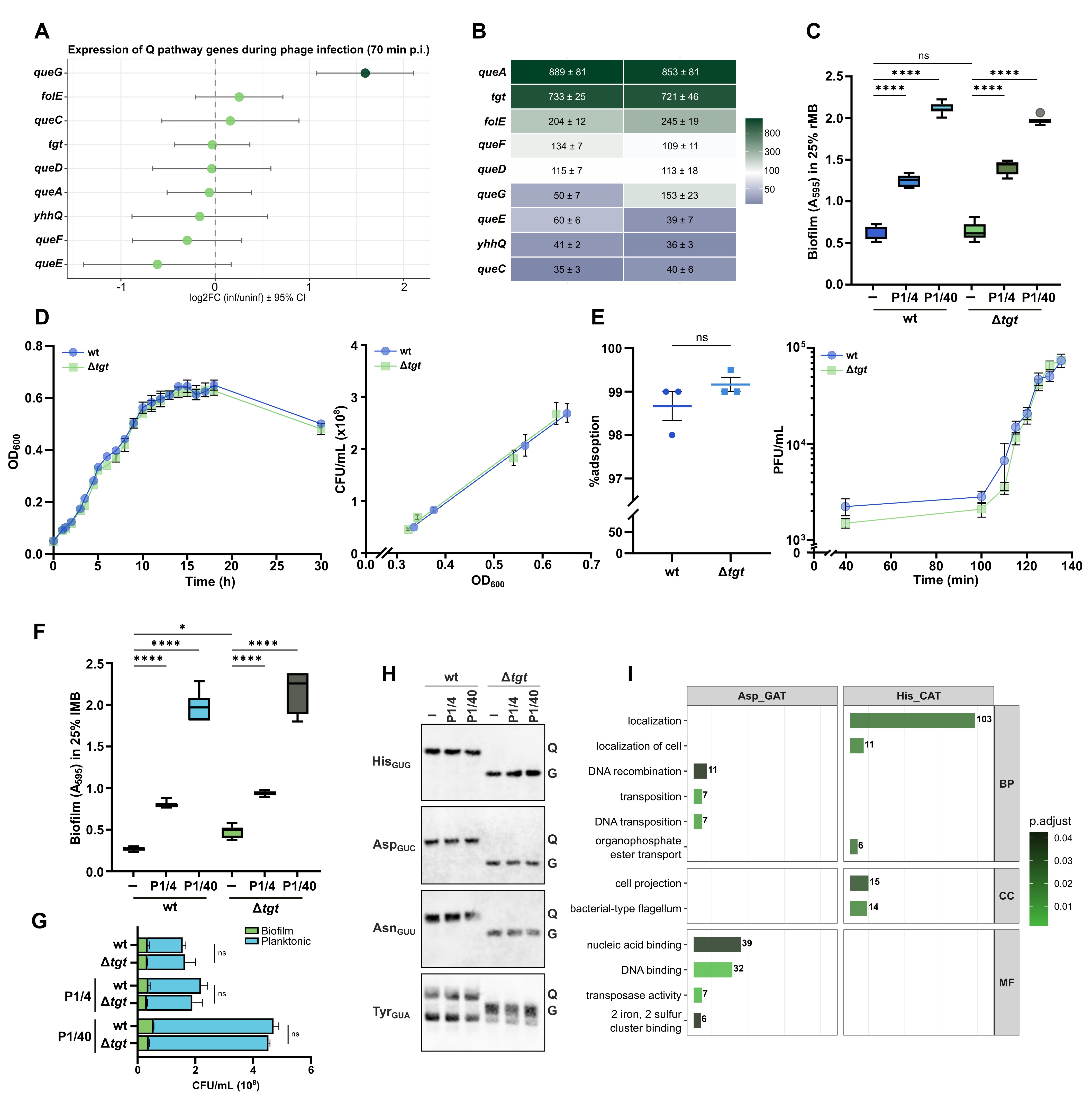

### Supplementary Figure 4

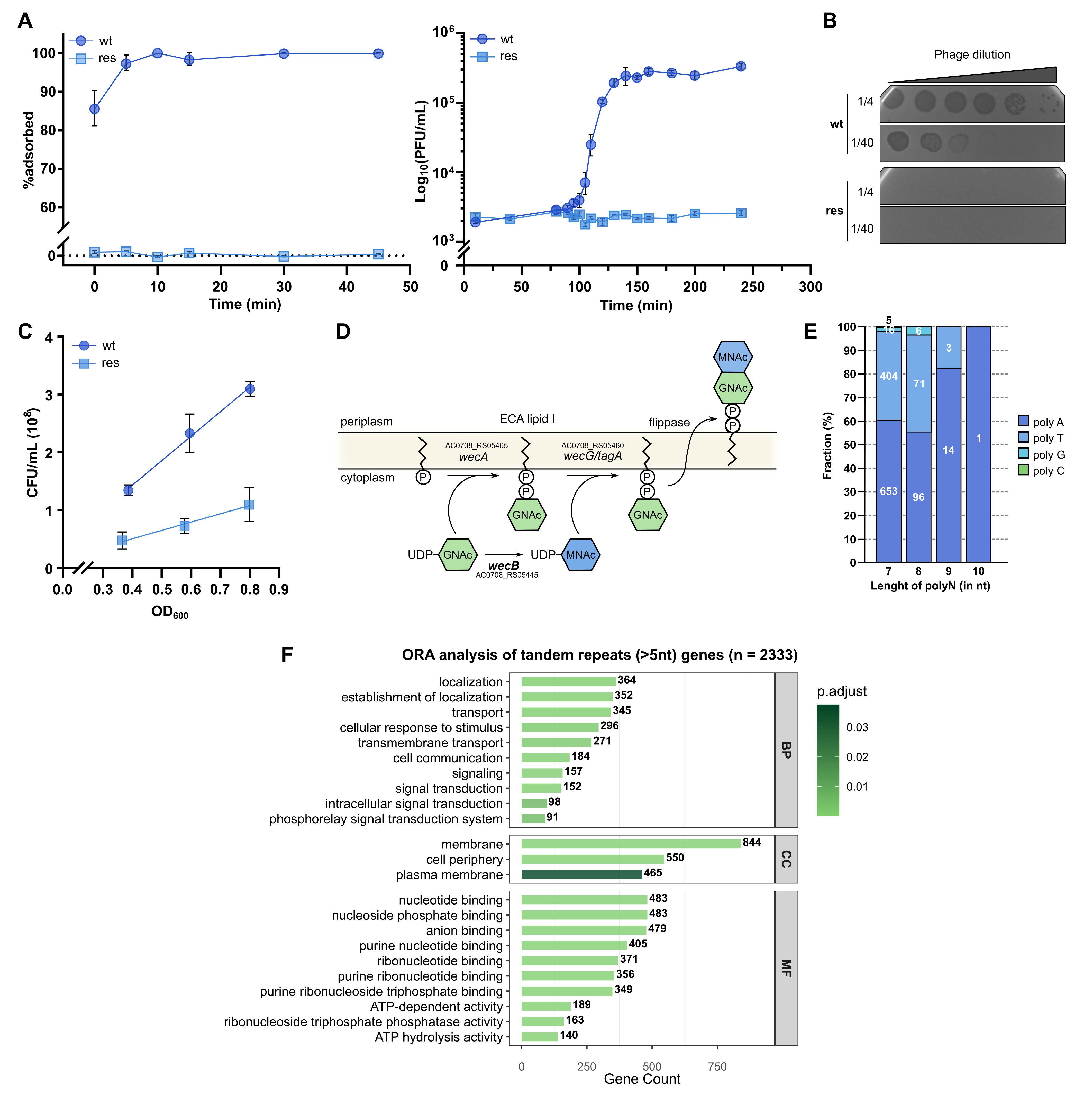

### Supplementary Figure 5

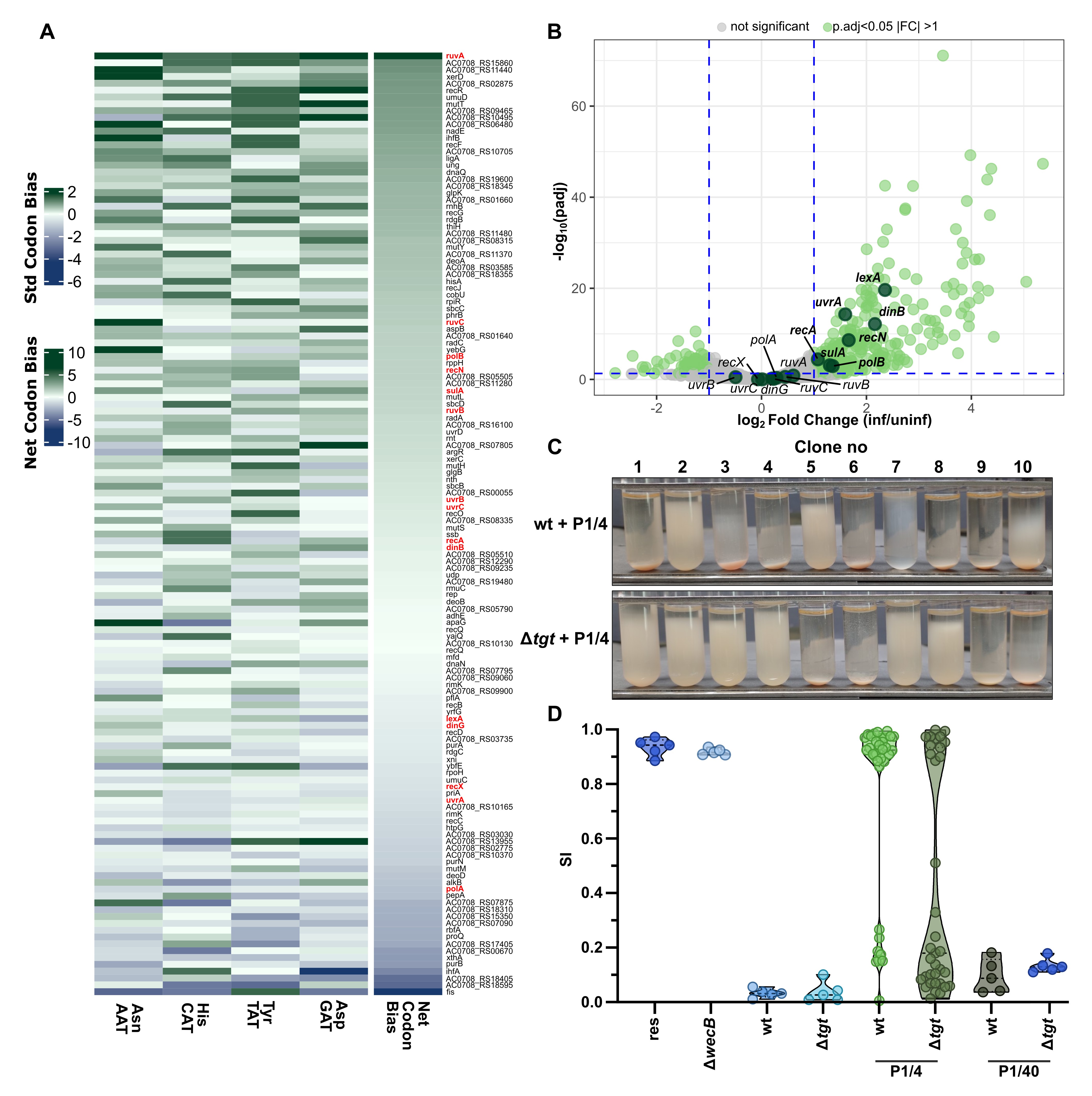

### Supplementary Figure 6

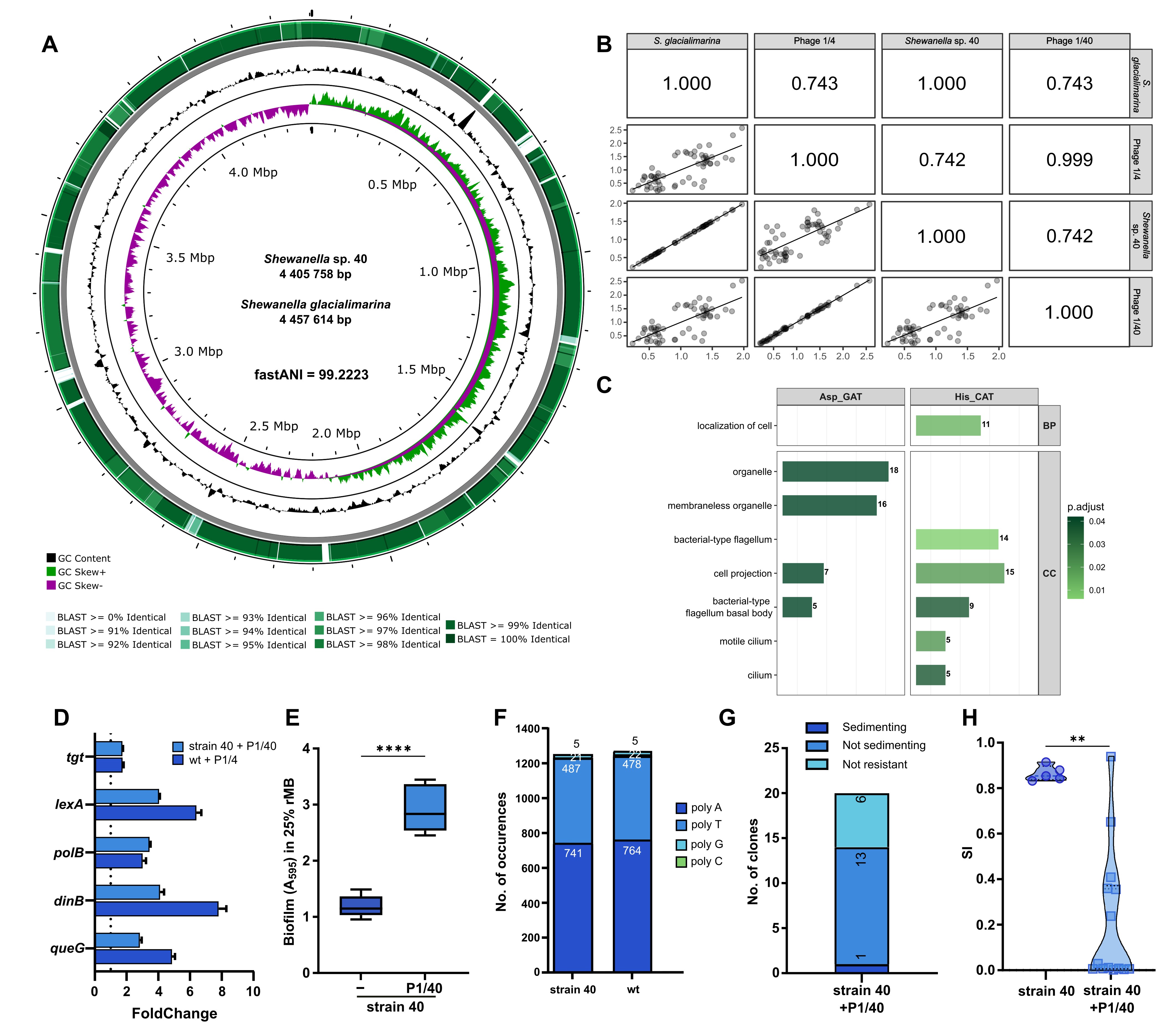

### Supplementary Figure 7

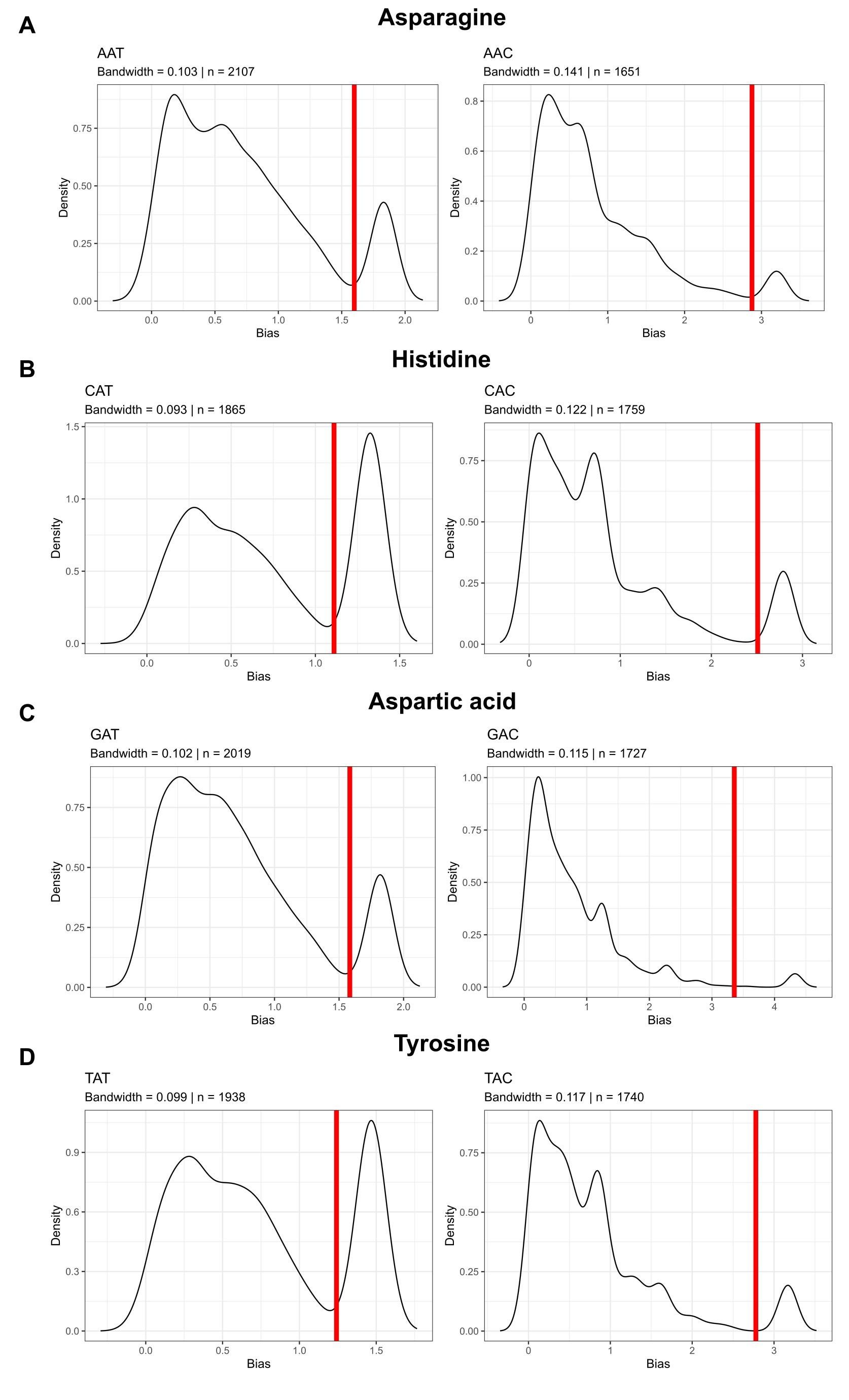

### Supplementary Figure 8

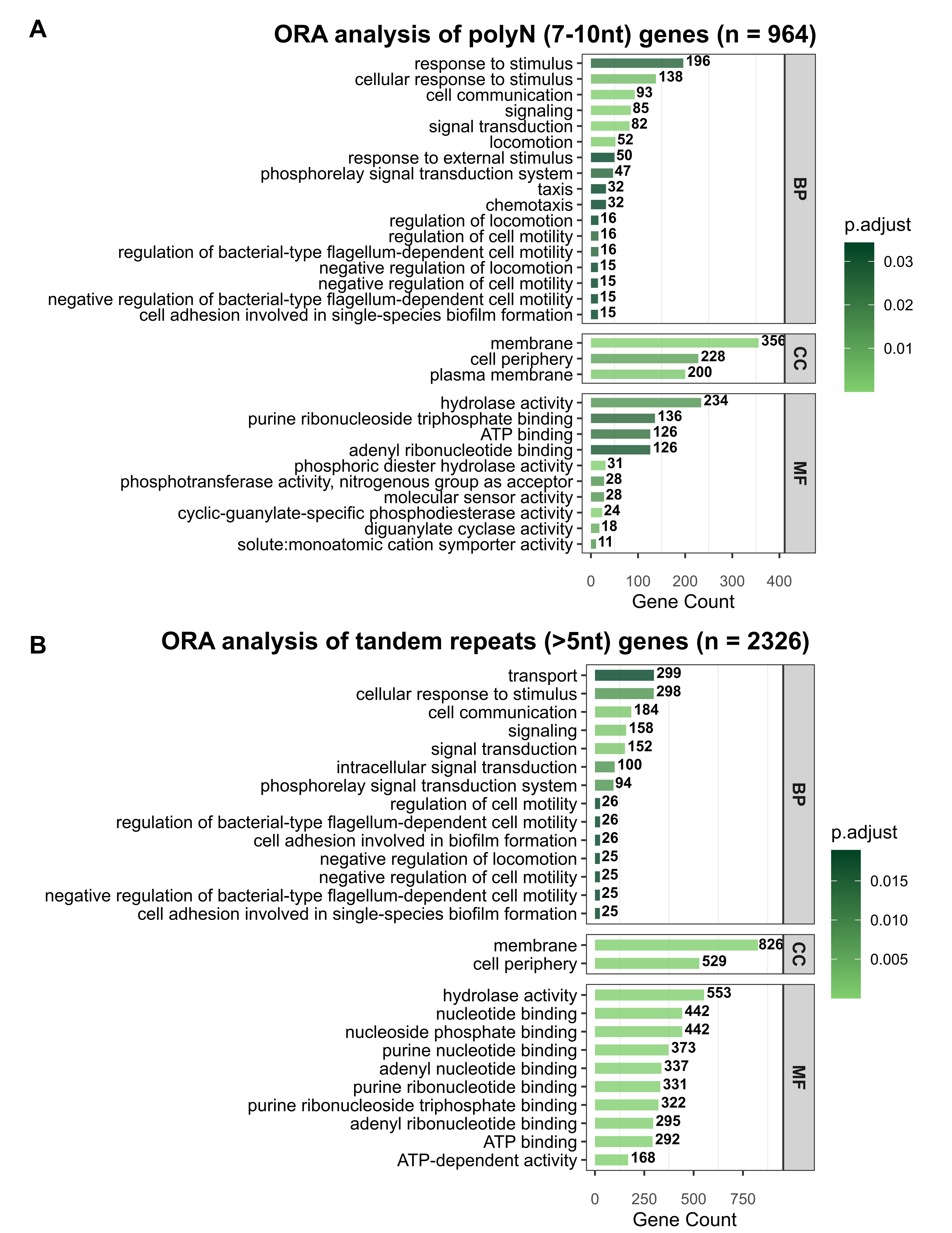
